# Nuclear Hormone Receptor NHR-49 controls a HIF-1-independent hypoxia adaptation pathway in *Caenorhabditis elegans*

**DOI:** 10.1101/2021.02.24.432575

**Authors:** Kelsie R. S. Doering, Xuanjin Cheng, Luke Milburn, Ramesh Ratnappan, Arjumand Ghazi, Dana L. Miller, Stefan Taubert

## Abstract

The response to insufficient oxygen (hypoxia) is orchestrated by the conserved Hypoxia-Inducible Factor (HIF). However, HIF-independent hypoxia response pathways exist that act in parallel to HIF to mediate the physiological hypoxia response. Here, we describe a HIF-independent hypoxia response pathway controlled by *Caenorhabditis elegans* Nuclear Hormone Receptor NHR-49, an orthologue of mammalian Peroxisome Proliferator-Activated Receptor alpha (PPARα). We show that *nhr-49* is required for worm survival in hypoxia and is synthetic lethal with *hif-1* in this context, demonstrating that these factors act independently. RNA-seq analysis shows that in hypoxia *nhr-49* regulates a set of genes that are *hif-1-*independent, including autophagy genes that promote hypoxia survival. We further show that Nuclear Hormone Receptor *nhr-67* is a negative regulator and Homeodomain-interacting Protein Kinase *hpk-1* is a positive regulator of the NHR-49 pathway. Together, our experiments define a new, essential hypoxia response pathway that acts in parallel to the well-known HIF-mediated hypoxia response.

## Introduction

Organisms are continuously exposed to endogenous and exogenous stresses, from suboptimal temperatures to foreign substances. Thus, an organism’s ability to mount specific stress responses, including protecting healthy cells from harm or inducing apoptosis when damage to a cell cannot be overcome, is critical for survival. Hypoxia is a stress that occurs when cellular oxygen levels are too low for normal physiological functions. It occurs naturally in cells and tissues during development, as well as in many diseases (P. Lee et al., 2020; Powell-Coffman, 2010). For example, due to hyperproliferation, inadequate vascularization, and loss of matrix attachment, cancer cells grow in hostile microenvironments featuring hypoxia. Certain cancers thus hijack the hypoxia response to allow growth and metastasis in these harsh conditions (Rankin & Giaccia, 2016; Schito & Semenza, 2016; T. Zhang et al., 2019), and tumor hypoxia correlates with poor clinical outcome (Keith & Simon, 2007). Most prominently, mutations in the tumor suppressor von Hippel Lindau (VHL), which inhibits the transcription factor hypoxia-inducible factor (HIF), occur in kidney cancers, and the resulting accumulation of HIF drives tumor growth (Kaelin, 2008; M. Li & Kim, 2011). In line with a pivotal role of HIF in these cancers are studies showing promising effects of HIF inhibitors in preclinical (Albadari et al., 2019; W. Chen et al., 2016; Cho et al., 2016) and clinical studies (Fallah & Rini, 2019). However, a better understanding of the transcriptional hypoxia adaptation pathway is needed to pinpoint new drug targets and to gain a deeper insight into how cells, tissues, and organisms cope with hypoxia.

The pathways that regulate the response to hypoxia are evolutionarily conserved from the nematode worm *Caenorhabditis elegans* to humans. As in mammals, a key pathway in *C. elegans* involves the transcription factor HIF-1, which is critical for the cellular responses to, and the defense against hypoxia (Choudhry & Harris, 2018; Jiang et al., 2001). To survive hypoxia, animals activate the EGL-Nine homolog (EGLN)–VHL-HIF pathway (*egl-9*–*vhl-1*–*hif-1* in *C. elegans*). In normoxic conditions (21% O_2_), HIF-1 is degraded and thus inactive. This occurs when EGL-9 adds a hydroxyl group onto a proline residue within HIF-1. The hydroxylated proline promotes binding of the E3 ubiquitin ligase VHL-1, leading to poly-ubiquitination and proteasomal degradation of HIF-1. However, in hypoxic conditions, EGL-9 is rendered inactive; hence, HIF-1 is stabilized and activates a hypoxia adaptation gene expression program (Epstein et al., 2001; Powell-Coffman, 2010).

Although the responses controlled by the HIF-1 master regulator are most studied, evidence for parallel transcriptional programs in hypoxia exists, from *C. elegans* to mammalian organisms. For example, the transcription factor B lymphocyte-induced maturation protein 1 (BLMP-1) has a *hif-1*-independent hypoxia regulatory role in *C. elegans* (Padmanabha et al., 2015), as does the conserved nuclear hormone receptor (NHR) estrogen-related receptor (dERR) in *Drosophila melanogaster* (Y. Li et al., 2013), and the cargo receptor Sequestosome 1 (SQSTM1/p62) in mammals (Pursiheimo et al., 2009). Thus, despite the evolutionarily conserved and important role of the HIF family, robust and effective hypoxia adaptation requires an intricate network of factors that act in concert. Compared to HIF, there is far less known about the mechanisms by which these pathways contribute to the hypoxia response.

*C. elegans* NHR-49 is a transcription factor orthologous to mammalian hepatocyte nuclear factor 4 (HNF4) and peroxisome proliferator-activated receptor α (PPARα) (K. Lee et al., 2016). Similar to these NHRs, it controls lipid metabolism by activating genes involved in fatty acid desaturation and mitochondrial β-oxidation (Pathare et al., 2012; Marc R. Van Gilst et al., 2005). By maintaining lipid homeostasis, NHR-49 is able to extend lifespan, a phenotype often associated with stress resistance (Burkewitz et al., 2015; Ratnappan et al., 2014). In addition to regulating lipid metabolism, NHR-49 also regulates putative xenobiotic detoxification genes in a dietary restriction-like state and during starvation (Chamoli et al., 2014; Goh et al., 2018), is required for resistance to oxidative stress (Goh et al., 2018), and activates innate immune response programs upon infection of *C. elegans* with *Staphylococcus aureus* (Wani et al., 2020), *Pseudomonas aeruginosa* (Naim et al., 2020), and *Enterococcus faecalis* (Dasgupta et al., 2020). Moreover, a recent report showed that *nhr-49* is required to increase expression of the Catechol-O-Methyl-Transferase *comt-5* downstream of the Hypoxia Inhibited Receptor tyrosine kinase *hir-1*, which mediates extracellular matrix remodelling in hypoxia (Vozdek et al., 2018). However, the role of *nhr-49* in hypoxia and how it intersects with *hif-1* have not been explored.

The detoxification gene flavin mono-oxygenase 2 (*fmo-2)* is induced in many of the aforementioned stresses in an *nhr-49*-dependent manner (Dasgupta et al., 2020; Goh et al., 2018; Wani et al., 2020). Interestingly, *fmo-2* is also a *hif-1*-dependent hypoxia response gene (Leiser et al., 2015; Shen et al., 2005), but its dependence on *nhr-49* in hypoxia is not known. We hypothesized that *nhr-49* may play a role in the worm hypoxia response, in part by regulating *fmo-2* expression. Here, we show that *nhr-49* is not only required to induce *fmo-2*, but controls a broad transcriptional response to hypoxia, including the induction of autophagy genes that are also required for survival in hypoxia. Our epistasis experiments indicate that *nhr-49* is functionally required independently of *hif-1* in hypoxia. Finally, we identify the protein kinase homeodomain interacting protein kinase 1 (*hpk-1*) as an upstream activator and the transcription factor *nhr-67* as a repressor of the *nhr-49* hypoxia response pathway. Together, our data define NHR-49 as a core player in a novel hypoxia response pathway that acts independently of *hif-1*.

## Results

### *nhr-49* is required to induce the expression of *fmo-2* in hypoxia

*C. elegans fmo-2* is induced by oxidative stress, starvation, and pathogen infection in an *nhr-49*-dependent fashion (Dasgupta et al., 2020; Goh et al., 2018; Wani et al., 2020). *fmo-2* expression is also induced in a *hif-1*-dependent manner during hypoxia (0.1% O_2_; Leiser et al., 2015; Shen et al., 2005). To test whether *nhr-49* regulates *fmo-2* expression in hypoxia, we quantified *fmo-2* mRNA levels in normoxia (21% O_2_) and hypoxia (0.5% O_2_) by quantitative Reverse Transcription PCR (qRT-PCR) in wild-type and mutant worms. The *nr2041* allele deletes portions of both the DNA binding domain and the ligand binding domain of *nhr-49* and is a predicted molecular null allele (Liu et al., 1999), and the *ia4* allele deletes exons 2-4 of *hif-1* and is also a predicted null allele (Jiang et al., 2001). In wild-type worms, *fmo-2* transcript levels increased approximately 40-fold in hypoxia, but this induction was blocked in both *nhr-49(nr2041)* and *hif-1(ia4)* mutant worms (Figure 1A). Experiments using a transgenic strain expressing a transcriptional *Pfmo-2::gfp* reporter (Goh et al., 2018) corroborated these observations *in vivo*. In normoxia, this reporter is weakly expressed in some neurons and in the intestine of transgenic animals, but expression was significantly elevated in the intestine of transgenic worms in hypoxia (Figure 1B-C). High pharyngeal expression made it difficult to quantify neuronal *Pfmo-2::gfp* in hypoxia. Consistent with our qRT-PCR data, loss of *nhr-49* abrogated the increase in intestinal upregulation of *Pfmo-2::gfp* worms following hypoxia exposure. We conclude that *nhr-49* is required to induce *fmo-2* in hypoxia.

**Figure 1.**
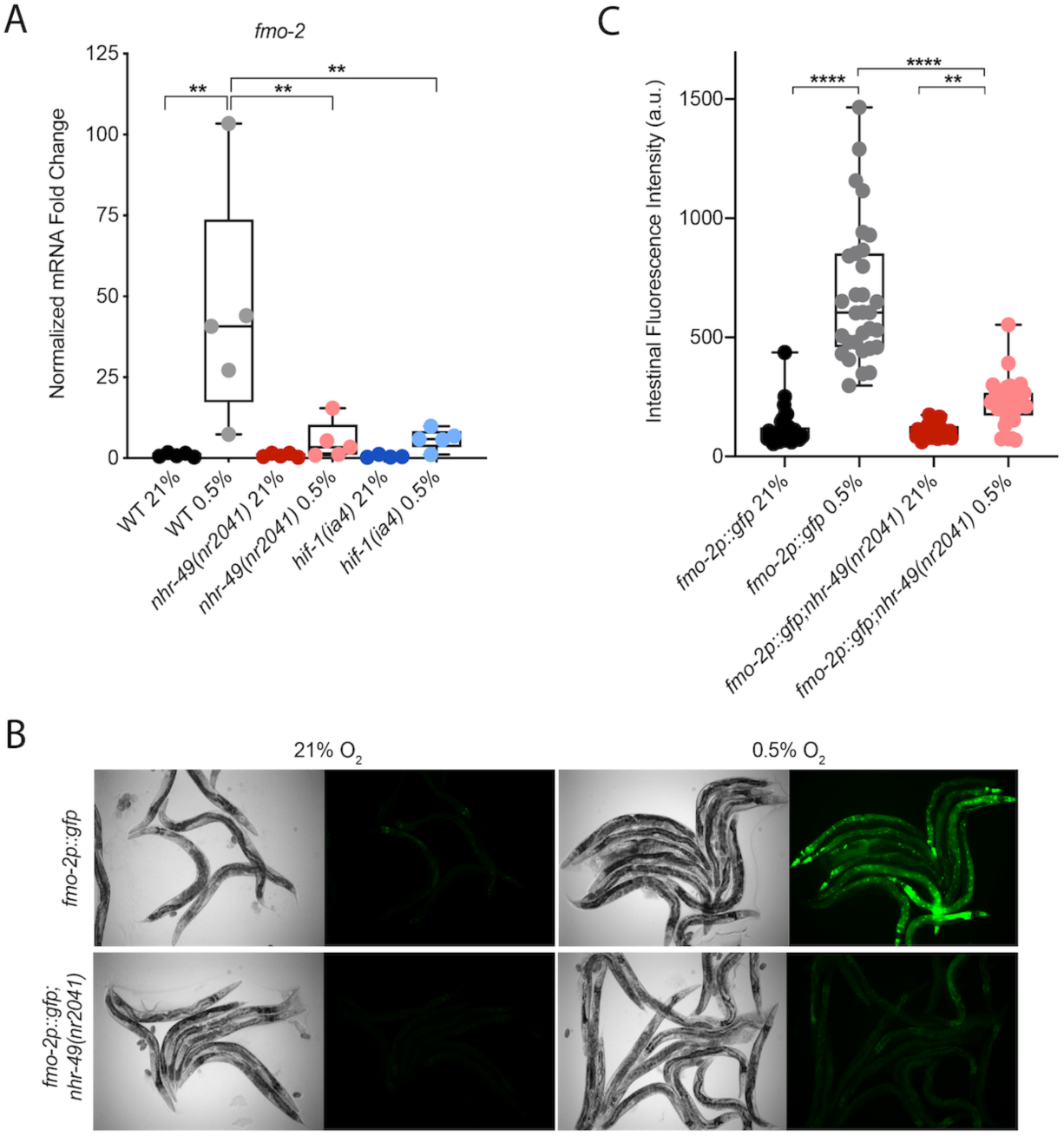
*nhr-49* regulates *fmo-2* induction following exposure to hypoxia. **(A)** The graph indicates fold changes of mRNA levels (relative to unexposed wild-type) in L4 wild-type, *nhr-49(nr2041),* and *hif-1(ia4)* worms exposed to room air (21% O_2_) or 0.5% O_2_ for 3 hr (n = 5). *,** p < 0.05, 0.01 (ordinary one-way ANOVA corrected for multiple comparisons using the Tukey method). **(B)** Representative micrographs show *Pfmo-2::gfp* and *Pfmo-2::gfp;nhr-49(nr2041)* adult worms in room air or following 4 hr exposure to 0.5% O_2_ and 1 hr recovery in 21% O_2_. **(C)** The graph shows the quantification of intestinal GFP levels in *Pfmo-2::gfp* and *Pfmo-2::gfp;nhr-49(nr2041)* worms following 4 hr exposure to 0.5% O_2_ and 1 hr recovery in 21% O_2_ (three repeats totalling >30 individual worms per genotype). **,**** p <0.01, 0.0001 (ordinary one-way ANOVA corrected for multiple comparisons using the Tukey method). WT = wild-type.

### *nhr-49* is required throughout the *C. elegans* life cycle to promote hypoxia resistance independently of *hif-1*

Wild-type embryos can survive a 24 h exposure to environments with as little as 0.5% O_2_, dependent on the presence of *hif-1* (Jiang et al., 2001; Nystul & Roth, 2004). We wanted to determine if *nhr-49*, like *hif-1*, is functionally required for worm survival during hypoxia. We first assessed the ability of worm embryos to survive for 24 hours in 0.5% O_2_ and then recover to the L4 or later stage when placed back in normoxia for 65 hours. We found that 86% of wild-type worm embryos reached at least the L4 stage, while only 25% of *nhr-49* and *hif-1* null mutant worms reached at least the L4 stage by that time (Figure 2A). This shows that, like *hif-1*, *nhr-49* is required for embryo survival in hypoxia.

**Figure 2.**
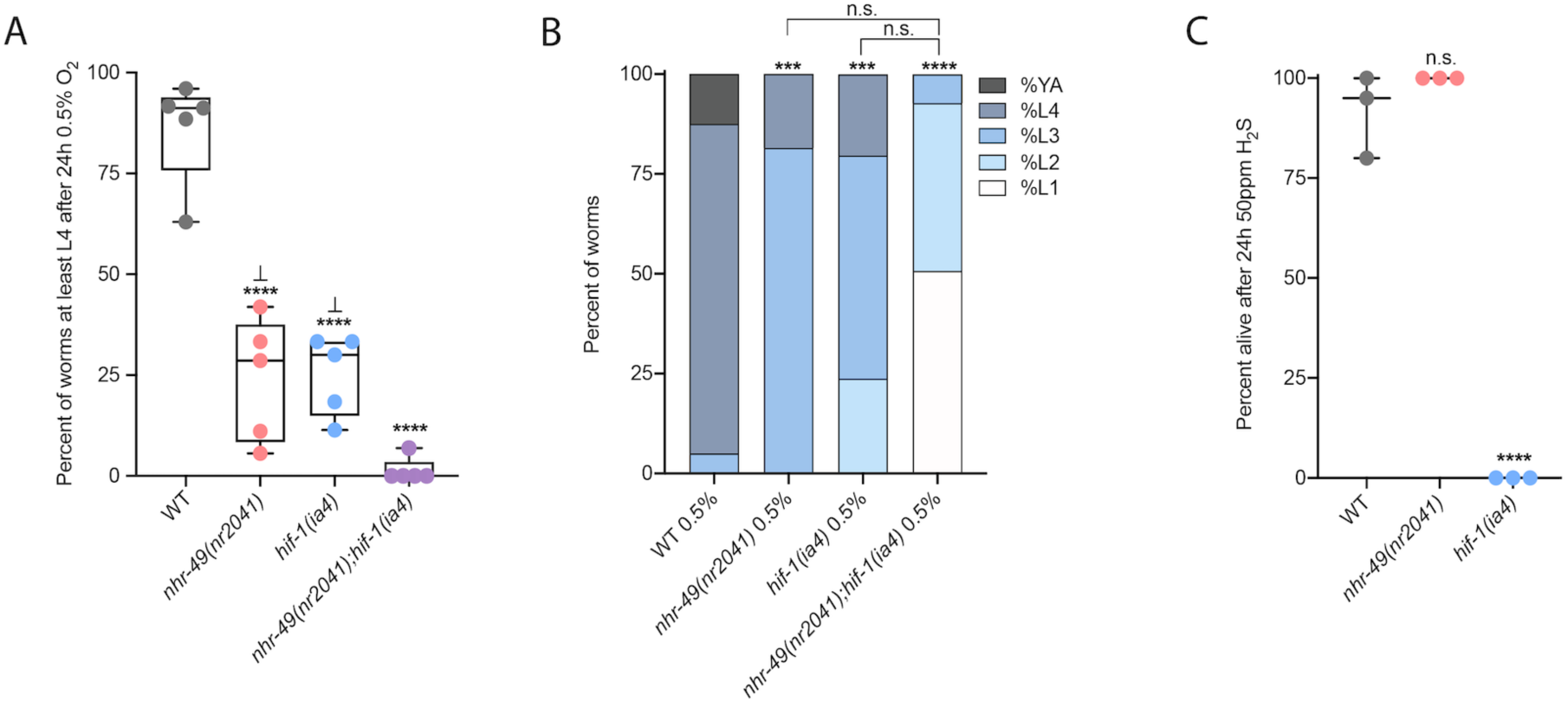
*nhr-49* and *hif-1* act in separate hypoxia response pathways at two stages of the worm life cycle. **(A)** The graph shows average population survival of wild-type, *nhr-49(nr2041)*, *hif-1(ia4)*, and *nhr-49(nr2041);hif-1(ia4)* worm embryos exposed for 24 hr to 0.5% O_2_ and then allowed to recover at 21% O_2_ for 65 hr, counted as ability to reach at least the L4 stage (five repeats totalling >100 individual worms per genotype). **** p<0.0001 vs. wild-type worms, ⊥ p<0.05 vs. *nhr-49(nr2041);hif-1(ia4)* (ordinary one-way ANOVA corrected for multiple comparisons using the Tukey method). **(B)** The graph shows average developmental success of wild-type, *nhr-49(nr2041)*, *hif-1(ia4)*, and *nhr-49(nr2041);hif-1(ia4)* larval worms following 48 hr exposure to 0.5% O_2_ from L1 stage (four repeats totalling >60 individual worms per genotype). ***, **** p<0.001, 0.0001 percent L4 or older vs. wild-type worms (ordinary one-way ANOVA corrected for multiple comparisons using the Tukey method). **(C)** The graph shows average population survival of wild-type, *nhr-49(nr2041)*, and *hif-1(ia4)* L4 worms following 24 hr exposure to 50 ppm hydrogen sulfide (three repeats totalling 60 individual worms per strain). **** p<0.0001 vs. wild-type worms (ordinary one-way ANOVA corrected for multiple comparisons using the Tukey method). n.s. = not significant, WT = wild-type.

Next, we asked whether *nhr-49* acts in the *hif-1* hypoxia response pathway or in a separate, parallel response pathway. To address this question, we generated a *nhr-49(nr2041);hif-1(ia4)* double null mutant. We observed that less than 2% of *nhr-49;hif-1* double null mutants reached at least the L4 stage following hypoxia exposure (Figure 2A). This suggests that *nhr-49* and *hif-1* act in separate, genetically independent hypoxia response pathways.

To determine if *nhr-49* and *hif-1* are required for larval development in hypoxia, we exposed newly hatched, first stage (L1) larvae to hypoxia for 48 hours. Following this treatment, 95% of wild-type worms reached at least the L4 stage (Figure 2B). In contrast, only 19% of *nhr-49* and only 20% of *hif-1* mutant worms, respectively, reached at least the L4 stage, and no *nhr-49;hif-1* double null mutant worms survived and developed to L4 (Figure 2B). Together, these results show that *nhr-49* is required for worm adaptation to hypoxia independently of *hif-1* both during embryogenesis and post-embryonically.

In normal conditions, *nhr-49* null worms have a shortened lifespan (Marc R. Van Gilst et al., 2005). This raised the concern that the defects observed in hypoxia may be an indirect consequence of NHR-49’s normal developmental roles. To test whether the effects observed above were due to a specific requirement for *nhr-49* in the hypoxia response, we studied worm development in normoxia. We found that loss of *nhr-49* had no effect on worm survival from embryo to at least the L4 stage at 21% O_2_ (Supplementary Figure 1A, Supplementary Table 1). Additionally, at 21% O_2_, *nhr-49* null mutants did not significantly develop slower than wildtype worms (Supplementary Figure 1B, Supplementary Table 2). Together, these data show that although *nhr-49* null mutants display mild developmental defects in normoxia, the phenotypes observed are due to the requirement for *nhr-49* specifically during hypoxia.

### *nhr-49* is dispensable for survival in hydrogen sulfide

To assess whether *nhr-49* is involved other responses requiring *hif-1*, we next asked if it was required for adaptation to hydrogen sulfide (H_2_S). H_2_S is produced endogenously and is an important signalling molecule in animals, including in *C. elegans* (L. Li et al., 2011). However, exposure to high levels of hydrogen sulfide can be lethal. As in the hypoxia response, *hif-1* is a master regulator of the transcriptional response to exogenous hydrogen sulfide, and *hif-1* is required for worm survival to 50ppm H_2_S. In (Budde & Roth, 2010; Miller et al., 2011)contrast, we found that *nhr-49* null mutants survive exposure to 50ppm H_2_S as well as wild-type controls (Figure 2C). This suggests that the requirement for *nhr-49* is stress specific, and that *nhr-49* does not participate in all *hif-1-*dependent stress responses. This is consistent with previous observations that the *hif-1*-dependent changes in gene expression in H_2_S are quite different than those seen in hypoxia (Miller et al., 2011). Additionally, the ability of *nhr-49* mutants to readily adapt to H_2_S provides further evidence that the mild developmental defects of *nhr-49* null mutants do not render the animal sensitive to all stresses. Instead, our data indicate that *nhr-49*’s requirement for hypoxia survival is due to a specific function for this regulator in this particular stress condition.

### The *nhr-49*-dependent transcriptional response to hypoxia includes *hif-1*-independent genes

To delineate the genes and biological processes regulated by NHR-49 in hypoxia, we analyzed whole-animal transcriptomes of wild-type, *nhr-49*, and *hif-1* worms before and after exposure to hypoxia using RNA-sequencing (RNA-seq; Figure 3A, B, Supplementary Figure 2A). We found that hypoxia significantly upregulated 718 genes and downregulated 339 genes more than two-fold in wild-type worms, including the known hypoxia-inducible genes *egl-9*, *phy-2*, *nhr-57*, and F22B5.4 (Bishop et al., 2004; Shen et al., 2005), validating our approach. 315 of the upregulated and 177 of the downregulated genes were dependent on *nhr-49*. Of these *nhr-49* regulated genes, 83 of the upregulated and 51 of the downregulated genes were *hif-1*-independent (Figure 3A, B). In line with our above data, *fmo-2* was induced in an *nhr-49*-dependent manner (Figure 3C). However, although our qRT-PCR data (Figure 1A) show that *fmo-2* induction is dependent on *hif-1*, our RNA-seq analysis excluded *fmo-2* from the *hif-1*-dependent set because it retained more than two-fold induction in hypoxia vs. normoxia (Supplementary Figure 2B). This suggests that although *fmo-2* induction is somewhat dependent on *hif-1*, it requires *nhr-49.* Thus, although many hypoxia responsive genes are controlled by both transcription factors, a subset is *nhr-49-*dependent but *hif-1*-independent.

**Figure 3.**
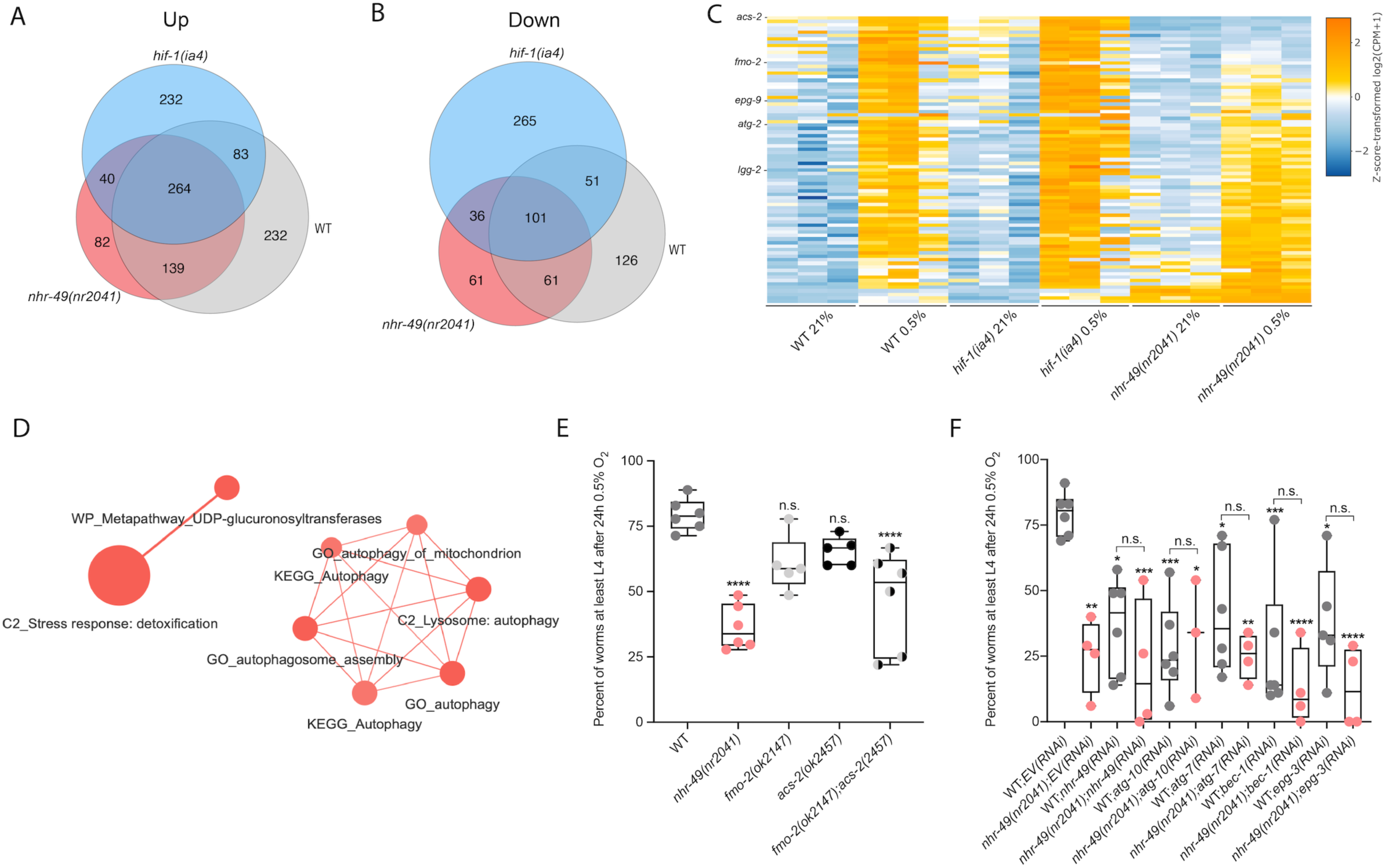
RNA-seq reveals an *nhr-49*-dependent transcriptional program in hypoxia. **(A, B)** Venn diagrams show the overlap of sets of hypoxia (0.5% O_2_; vs. normoxia 21% O_2_) regulated genes identified in differential expression analysis comparing wild-type, *nhr-49(nr2041)*, and *hif-1(ia4)* worms. Numbers indicate the number of significantly differentially expressed genes that are upregulated (A) and downregulated (B) at least two-fold. **(C)** Heatmap of the expression levels of the 83 genes which are significantly induced over two-fold in 21% O_2_ vs. 0.5% O_2_ in wild-type and *hif-1(ia4)* worms, but not in *nhr-49(nr2041)*, i.e. *nhr-49*-dependent hypoxia response genes. Genes along the y-axis are colored in each repeat based on their z-scores of the log2-transformed Counts Per Million (CPM) plus 1. Notable genes are highlighted. **(D)** Network view of the enriched functional categories among the 83 genes, which are significantly induced over two-fold in 21% O_2_ vs. 0.5% O_2_ in wild-type and *hif-1(ia4)* worms, but not in *nhr-49(nr2041)*. Edges represent significant gene overlap as defined by a Jaccard Coefficient larger than or equal to 25%. The dot size reflects the number of genes in each functional category; colour intensity reflects statistical significance (−log10 p-value). **(E)** The graph shows the average population survival of wild-type, *nhr-49(nr2041)*, *fmo-2(ok2147)*, *acs-2(ok2457)*, and *fmo-2(ok2147);acs-2(ok2457)* worm embryos following 24 hr exposure to 0.5% O_2_, then allowed to recover at 21% O_2_ for 65 hr, and counted as ability to reach at least L4 stage (five or more repeats totalling >100 individual worms per strain). **** p<0.0001 vs. wild-type worms. Comparison of single mutants to *fmo-2(ok2147);acs-2(ok2457)* not significant (ordinary one-way ANOVA corrected for multiple comparisons using the Tukey method). **(F)** The graph shows the average population survival of second generation wild-type and *nhr-49(nr2041)* worm embryos fed *EV*, *nhr-49*, *atg-10*, *atg-7*, *bec-1*, or *epg-3* RNAi, followed by 24 hr exposure to 0.5% O_2_ and recovery at 21% O_2_ for 65 hr, and counted as ability to reach at least L4 stage (three or more repeats totalling >100 individual worms per strain). *, **, ***, **** p<0.05, 0.01, 0.001, 0.0001 vs. followed by worms fed *EV(RNAi)* (ordinary one-way ANOVA corrected for multiple comparisons using the Tukey method). n.s. = not significant, WT = wild-type.

We next performed functional enrichment analysis to identify the biological pathways and processes regulated in hypoxia, and specifically those dependent on *nhr-49.* In wild-type worms, pathways such as detoxification, response to heavy metal stress, and autophagy were induced (Supplementary Figure 2C), whereas processes such as amino acid transport and metabolism were downregulated (Supplementary Figure 2D). In the set of 83 genes that exclusively require *nhr-49* but not *hif-1* for induction in hypoxia (Figure 3C, Supplementary Table 3), autophagy and detoxification were significantly enriched (Figure 3D), suggesting a requirement for *nhr-49* to regulate these particular processes in hypoxia. Interestingly, a separate set of detoxification genes was dependent only on *hif-1* (Supplementary Figure 2E, F, Supplementary Table 4), and a third set of detoxification genes were independent of both *nhr-49* and *hif-1* (Supplementary Figure 2G, H, Supplementary Table 5). This suggests that there may be an additional transcription factor(s) regulating this process in hypoxia.

Our RNA-seq data revealed that the acyl-CoA synthetase gene *acs-2* is induced in response to hypoxia in an *nhr-49*-dependent manner (Figure 3C, Supplementary Table 3). ACS-2 acts in the first step of mitochondrial fatty acid β-oxidation, and is strongly induced by NHR-49 during starvation and following exposure to *E. faecalis* (Dasgupta et al., 2020; Marc R. Van Gilst et al., 2005). To validate our RNA-seq data, we quantified *acs-2* expression via qRT-PCR. Following hypoxia exposure, *acs-2* transcript levels increased approximately seven-fold, and this induction was blocked in the *nhr-49* null mutant, but not the *hif-1* null mutant (Supplementary Figure 3A). We used a transgenic strain expressing a transcriptional *Pacs-2::gfp* reporter to study this regulation i*n vivo* (Burkewitz et al., 2015). This reporter showed moderate GFP expression in the body of animals under normoxia, but expression increased substantially in the intestine following exposure to hypoxia (Supplementary Figure 3B, C). Consistent with our RNA-seq and qRT-PCR data, loss of *nhr-49* blocked transcriptional activation via the *acs-2* promoter, as GFP was weaker in the intestines of these worms following hypoxia exposure (Supplementary Figure 3B, C). Collectively, these data show that *nhr-49* is specifically required and that *hif-1* is dispensable for induction of *acs-2* in hypoxia.

### Autophagy genes are critical downstream targets of *nhr-49* in hypoxia

Next, we wanted to determine which of *nhr-49*’s downstream transcriptional targets are functionally important for worm survival in hypoxia. We first assessed the ability of *fmo-2(ok2147)* and *acs-2(ok2457)* embryos to survive hypoxia, as both genes are strongly induced during hypoxia in an *nhr-49*-dependent manner. Individually, loss of either *fmo-2* (60%) or *acs-2* (65%) did not significantly decrease embryo viability compared to wild-type (79%) (Figure 3E). However, simultaneous loss of both *fmo-2* and *acs-2* resulted in a significant decrease in survival after hypoxia (47%). None of the mutant animals showed embryo viability defects in normoxia, indicating that the phenotypes observed were specifically due to the requirement of these genes in hypoxia survival (Supplementary Figure 4A, Supplementary Table 1). These data suggest that *fmo-2* and *acs-2* each contribute only modestly to worm survival to hypoxia, and are likely not the main factors contributing to *nhr-49*’s importance in survival to this stress. This resembles previous observations that mutations that disrupt individual *hif-1*-responsive genes show only minor defects in hypoxia survival (Shen et al., 2005).

Our RNA-seq analysis revealed autophagy as a major biological process modulated by *nhr-49*. Notably, *C. elegans* show sensitivity to anoxia when the autophagy pathway is disrupted (Samokhvalov et al., 2008), and autophagy is upregulated in anoxia (Chapin et al., 2015). However, the responses to anoxia and hypoxia are mediated by different regulatory pathways (Nystul & Roth, 2004), and it thus was not *a priori* clear whether autophagy is also required for hypoxia resistance. To determine if upregulation of autophagy by *nhr-49* is required for worm survival in hypoxia, we depleted several autophagy genes using feeding RNA interference (RNAi) in the wild-type and *nhr-49* null mutant backgrounds and assessed the ability of these embryos to survive hypoxia. RNAi mediated knockdown of the autophagy genes *atg-10* (28%), *atg-7* (41%), *bec-1* (27%), and *epg-3* (38%) caused significant sensitivity to hypoxia in the wild-type background compared to the empty vector (EV) control RNAi treatment (79%; Figure 3F). Importantly, the sensitivity of worms did not change when these genes were knocked down in the *nhr-49* null background (32%, 25%, 13%, 13%, respectively), suggesting that these genes act in the same pathway as *nhr-49*. Depletion of these genes by RNAi alone did not cause impaired development from embryo to L4 in normoxia, indicating the phenotypes observed were specifically due to the requirement of these genes in hypoxia survival (Supplementary Figure 4B, Supplementary Table 1). Together, these data show that autophagy is a functionally important *nhr-49* regulated process required for worm survival in hypoxia.

### NHR-49 expression in multiple tissues is sufficient to promote hypoxia survival

To test if *nhr-49* activation is sufficient to promote survival of worms in hypoxia, we studied the *nhr-49(et13)* gain-of-function strain, which is sufficient to induce *fmo-2* (Goh et al., 2018; K. Lee et al., 2016). After 24 hours of exposure to hypoxia, approximately 86% of wild-type eggs develop to at least L4 stage (Figure 2A), but after 48 hours of hypoxia exposure, only approximately 44% of wild-type eggs develop to at least L4 stage (Figure 4A). In contrast, 75% of *nhr-49(et13)* gain-of-function eggs develop to at least L4 stage after 48 hours of hypoxia exposure, indicating that NHR-49 activation is sufficient to improve the population survival of worms in hypoxia.

**Figure 4.**
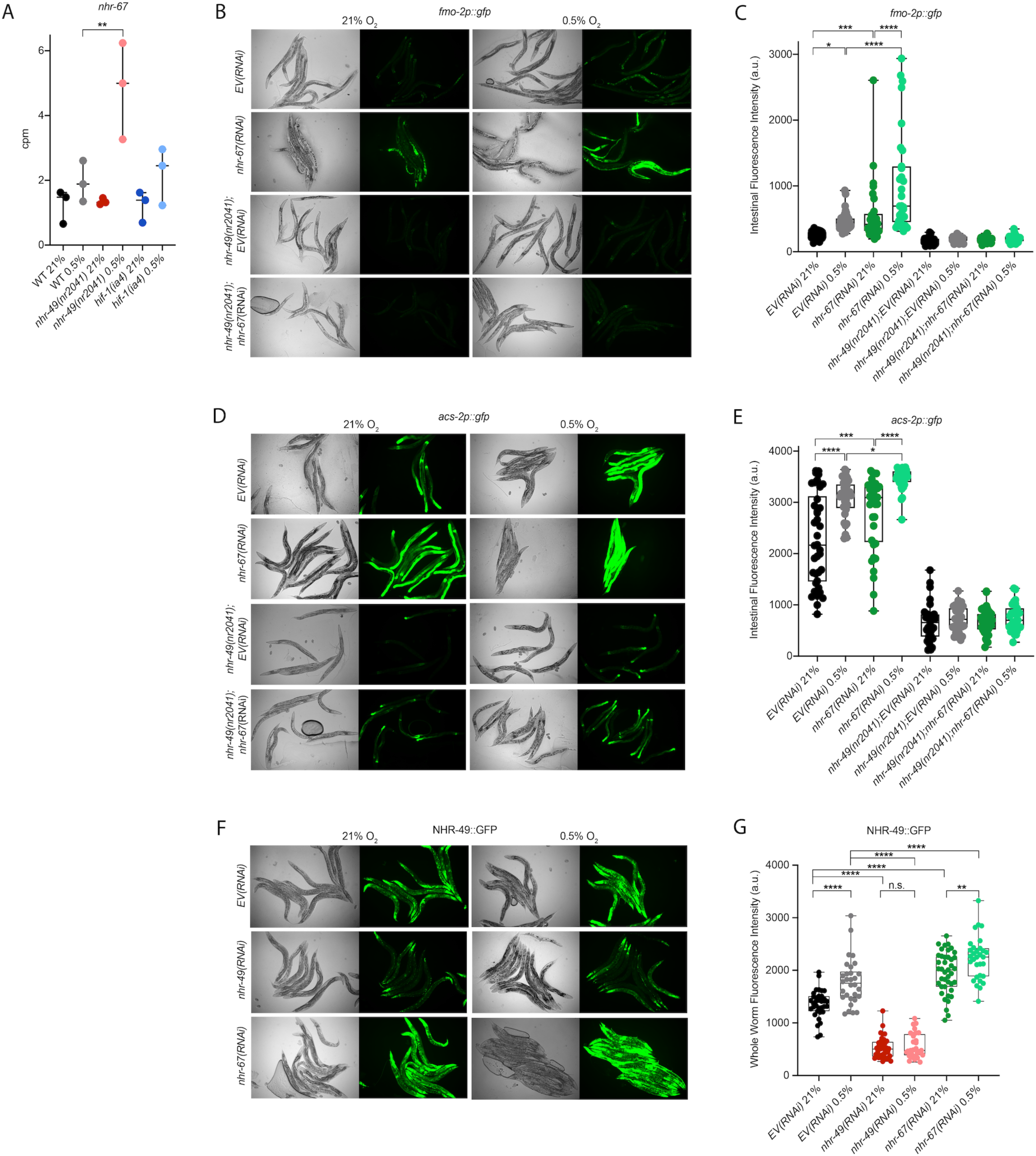
*nhr-49* is sufficient to promote survival in hypoxia and induce some hypoxia response genes. **(A)** The graph shows the average population survival of wild-type, *nhr-49(nr2041)*, and *nhr-49(et13)* worm embryos following 48 hr exposure to 0.5% O_2_, then allowed to recover at 21% O_2_ for 42 hr, and counted as ability to reach at least L4 stage (five repeats totalling >100 individual worms per strain). * p<0.05 vs. wild-type worms, ⊥⊥⊥ p<0.001 vs. *nhr-49(et13)* worms (ordinary one-way ANOVA corrected for multiple comparisons using the Tukey method). **(B)** The graph shows average population survival of *nhr-49* tissue specific rescue worm embryos following 24 hr exposure to 0.5% O_2_, then allowed to recover at 21% O_2_ for 65 hr, and counted as ability to reach at least L4 stage. *Pglp-19::nhr-49::gfp* for intestine, *Pcol-12::nhr-49::gfp* for hypodermis, *Prgef-1::nhr-49::gfp* for neurons, and *Pnhr-49::nhr-49::gfp* for endogenous (four or more repeats totalling >50 individual worms per strain). * p<0.05 vs. matching non-GFP siblings. **(C)** The graph shows fold changes of mRNA levels (relative to wild-type) in L4 *nhr-49(et13)* worms (n = 3). *,*** p < 0.05, 0.001 vs. wild-type worms (ordinary one-way ANOVA corrected for multiple comparisons using the Tukey method). WT = wild-type.

NHR-49 is expressed in multiple tissues, including the intestine, neurons, muscle, and hypodermis (Ratnappan et al., 2014). Neuronal NHR-49 is sufficient to extend life span in some contexts and regulates genes in distal tissues (Burkewitz et al., 2015), but where the protein acts to regulate the response to hypoxia is unknown. To study this, we induced expression of an NHR-49::GFP translational fusion protein in the *nhr-49(nr2041)* mutant background using tissue-specific promoters (Naim et al., 2020). Comparing the NHR-49::GFP rescue strains to their non-GFP siblings, we found that expressing *nhr-49* in the intestine, neurons, hypodermis, or from its endogenous promoter was sufficient to restore population survival to wild-type levels (Figure 4B). This suggests that NHR-49 can act in multiple somatic tissues to regulate the organismal hypoxia response.

To determine if NHR-49 activity alone is sufficient to induce expression of hypoxia response genes, we assessed the ability of the *nhr-49(et13)* gain-of-function strain to induce some of the *nhr-49-* dependent hypoxia response genes from our RNA-seq analysis in the absence of stress (Figure 4C). In line with previous findings (Goh et al., 2018; K. Lee et al., 2016), *nhr-49* was sufficient to induce *fmo-2* and *acs-2* expression on its own. However, other hypoxia inducible *nhr-49* regulated genes involved in autophagy and detoxification (Supplementary Table 3) were not induced in the *nhr-49(et13)* gain-of-function mutant. It is possible that *nhr-49* regulates autophagy indirectly in a manner independent of transcription, or that this *et13* mutation, which has combined gain and loss of function properties (K. Lee et al., 2016), cannot induce these tested autophagy genes. It is also possible that, to induce these genes, NHR-49 acts in concert with another hypoxia-responsive transcription factor, which is not activated in the *nhr-49(et13)* mutant. Together, this shows that NHR-49 is sufficient to extend survival of worms in hypoxia in various tissues, but the gain-of-function strain is only able to induce certain response genes without the presence of stress.

### The nuclear hormone receptor NHR-67 negatively regulates the *nhr-49* hypoxia response

Cellular stress response pathways are intricate networks involving a multitude of proteins. Activation or repression of downstream response genes thus often requires signaling via additional factors such as kinases and transcription factors. To identify additional factors acting in the *nhr-49* regulated hypoxia response pathway, we studied proteins that have been reported to physically interact with NHR-49 (Reece-Hoyes et al., 2013). One such interaction partner is the nuclear hormone receptor NHR-67, the sole *C. elegans* ortholog of the *D. melanogaster tailless* and vertebrate *NR2E1* genes (Gissendanner et al., 2004). NHR-67 is important in neural and uterine development (Fernandes & Sternberg, 2007; Verghese et al., 2011), but a role for this NHR in stress responses has not yet been described. Our RNA-seq data showed that *nhr-67* mRNA expression is modestly increased during hypoxia in wild-type worms, and much more substantially induced in the *nhr-49* null background (Figure 5A), suggesting a possible regulatory interaction between these two NHRs during hypoxia. To explore this interaction further, we used feeding RNAi to knock down *nhr-67* in normoxia and hypoxia, and observed how this affected the expression of the *Pfmo-2::gfp* and *Pacs-2::gfp* transcriptional reporters. Compared to *EV(RNAi),* knockdown of *nhr-67* significantly induced both reporters even in the absence of stress, suggesting a repressive role for *nhr-67* on these genes (Figure 5B-E). In hypoxia, *nhr-67(RNAi)* resulted in even higher expression of these reporters. In both normoxia and hypoxia, increased expression of the reporters was dependent on *nhr-49*, as loss of *nhr-49* abrogated the GFP induction (Figure 5B-E). The *nhr-49(et13)* gain-of-function mutation is sufficient to induce expression of the *Pfmo-2::gfp* reporter in non-stressed conditions (Goh et al., 2018), although it does not alter *nhr-67* expression under normoxic conditions (Supplementary Figure 5A). Knockdown of *nhr-67* further increased the expression of the *Pfmo-2::gfp* reporter in the *nhr-49(et13)* background in both normoxia and hypoxia (Supplementary Figure 5B, C). Together, these data suggest that *nhr-67* negatively regulates the expression of the hypoxia response genes *fmo-2* and *acs-2* in both normoxic and hypoxic conditions, and that this regulation is dependent on *nhr-49*.

**Figure 5.**
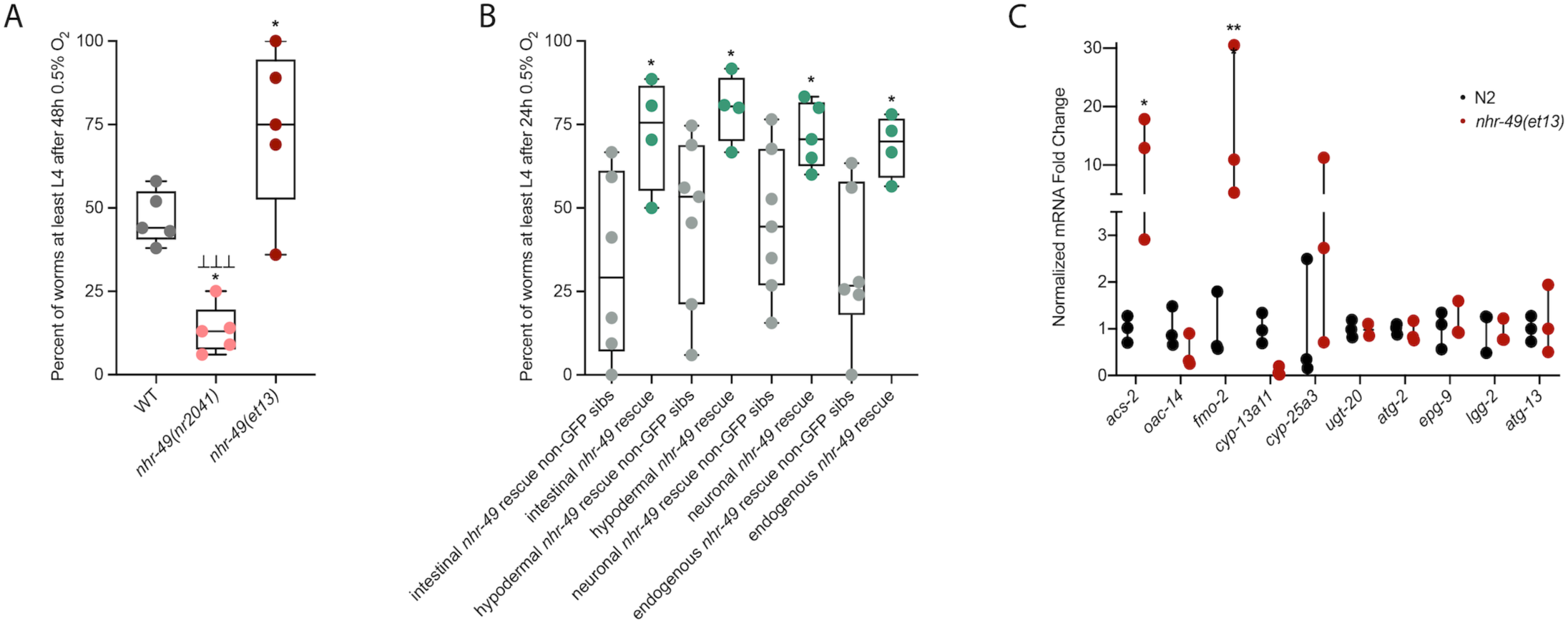
*nhr-67* is a negative regulator of the *nhr-49*-dependent hypoxia response pathway. **(A)** The graph shows average transcript levels in counts per million (CPM) of *nhr-67* mRNA in L4 wild-type, *nhr-49(nr2041)*, and *hif-1(ia4)* worms exposed to 0.5% O_2_ for 3 hr or kept at 21% O_2_ (n = 3). ** p <0.01 (ordinary one-way ANOVA corrected for multiple comparisons using the Tukey method). **(B-E)** Representative micrographs and quantification of intestinal GFP levels in *Pfmo-2::gfp* and *Pfmo-2::gfp;nhr-49(nr2041)* (B, C) and *Pacs-2::gfp* and *Pacs-2::gfp;nhr-49(nr2041)* (D, E) adult worms fed EV RNAi or *nhr-67* RNAi following 4 hr exposure to 0.5% O_2_ and 1 hr recovery in 21% O_2_ (three repeats totalling >30 individual worms per strain). *,***,**** p <0.05, 0.001, 0.0001 (ordinary one-way ANOVA corrected for multiple comparisons using the Tukey method). **(F)** Representative micrographs show *Pnhr-49::nhr-49::gfp* adult worms fed EV, *nhr-49*, or *nhr-67* RNAi following 4 hr exposure to 0.5% O_2_ and 1 hr recovery in 21% O_2_. **(G)** The graph shows quantification of whole worm GFP levels in *Pnhr-49::nhr-49::gfp* worms fed EV, *nhr-49*, or *nhr-67* RNAi following 4 hr exposure to 0.5% O_2_ and 1 hr recovery in 21% O_2_ (three or more repeats totalling >30 individual worms per strain). **** p <0.0001 (ordinary one-way ANOVA corrected for multiple comparisons using the Tukey method). n.s. = not significant, WT = wild-type.

As a negative regulator of *nhr-49*-dependent hypoxia response genes, it is possible that *nhr-67* acts upstream of *nhr-49* or directly on the promoter of *acs-2* and *fmo-2*. To determine how *nhr-67* regulates this response, we used feeding RNAi to knock down *nhr-67* and observed expression of the *Pnhr-49::nhr-49::gfp* translational reporter (which encodes GFP tagged to a full length NHR-49 transgene under control of its endogenous promoter from extra-chromosomal arrays, henceforth referred to as NHR-49::GFP; Ratnappan et al., 2014). The NHR-49::GFP protein is expressed most highly in the intestine, and also shows expression in neurons, muscle, and the hypodermis (Ratnappan et al., 2014). Whole worm NHR-49::GFP expression was increased in both normoxia and hypoxia following knockdown of *nhr-67*, with the highest increase observed in the intestine (Figure 5F, G). This suggests that *nhr-67* negatively regulates NHR-49, but in hypoxia, an increase in NHR-49 protein levels may in turn repress *nhr-67*, suggesting a negative feedback loop. The effects seen on *fmo-2* and *acs-2* expression are likely a consequence of NHR-67’s effect on NHR-49.

Loss of function mutations in *nhr-67* cause early L1 lethality or arrest (Fernandes & Sternberg, 2007), so we used feeding RNAi to study *nhr-67*’s functional requirements in hypoxia. We assessed the ability of *nhr-67(RNAi)* embryos to survive hypoxia and recover, as described above. Resembling *nhr-49(RNAi)* worms (46% survival), only 49% of *nhr-67* knockdown embryos survived to at least L4 stage compared to the *EV(RNAi)* worms (73%; Supplementary Figure 5D). Although RNAi knockdown of *nhr-67* and *nhr-49* causes developmental delays, the majority of *nhr-67(RNAi*) and *nhr-49(RNAi*) worms were able to reach at least L4 stage in normoxia (91% and 90%, respectively), resembling *EV(RNAi)* worms (100%; Supplementary Figure 5E, Supplementary Table 1). Thus, although *nhr-67* appears to perform a negative regulatory role on the hypoxia pathway, it, too, is functionally required for survival in hypoxia. Taken together, these data show that *nhr-67* is a functionally important negative regulator of the *nhr-49-*dependent hypoxia response.

### The kinase *hpk-1* positively regulates *nhr-49*-dependent hypoxia response genes and is required for survival in hypoxia

To identify additional factors acting in the *nhr-49*-dependent hypoxia response pathway, we studied eight kinases that we found to potentially act in the *nhr-49*-dependent oxidative stress response (Doering & Taubert, manuscript in preparation). We depleted each kinase using feeding RNAi to determine if any treatment prevented *Pfmo-2::gfp* induction in hypoxia in the worm intestine. As expected, *nhr-49* RNAi diminished this intestinal fluorescence compared to the *EV(RNAi)* (Figure 6A, B). Of the eight kinases tested, RNAi knockdown of the nuclear serine/threonine kinase homeodomain interacting protein kinase 1 (*hpk-1*) significantly decreased intestinal *Pfmo-2::gfp* expression following hypoxia exposure (Figure 6A, B), phenocopying *nhr-49* knockdown. Knockdown of *hpk-1* also significantly reduced intestinal expression of the *Pacs-2::gfp* reporter in hypoxia (Figure 6C, D) and reduced expression of *Pfmo-2::gfp* in the *nhr-49(et13)* background (Supplementary Figure 6A, B). We corroborated these data using qRT-PCR in wild-type worms and a *hpk-1(pk1393)* mutant. The *pk1393* allele deletes the majority of the kinase domain of *hpk-1* and is a predicted molecular null allele (Raich et al., 2003). In hypoxia, the expression of both *acs-2* and *fmo-2* was significantly reduced by loss of *hpk-1*, phenocopying loss of *nhr-49* (Figure 6E). Together, these data suggest that, like *nhr-49*, *hpk-1* is required for upregulation of *fmo-2* and *acs-2* in response to hypoxia.

**Figure 6.**
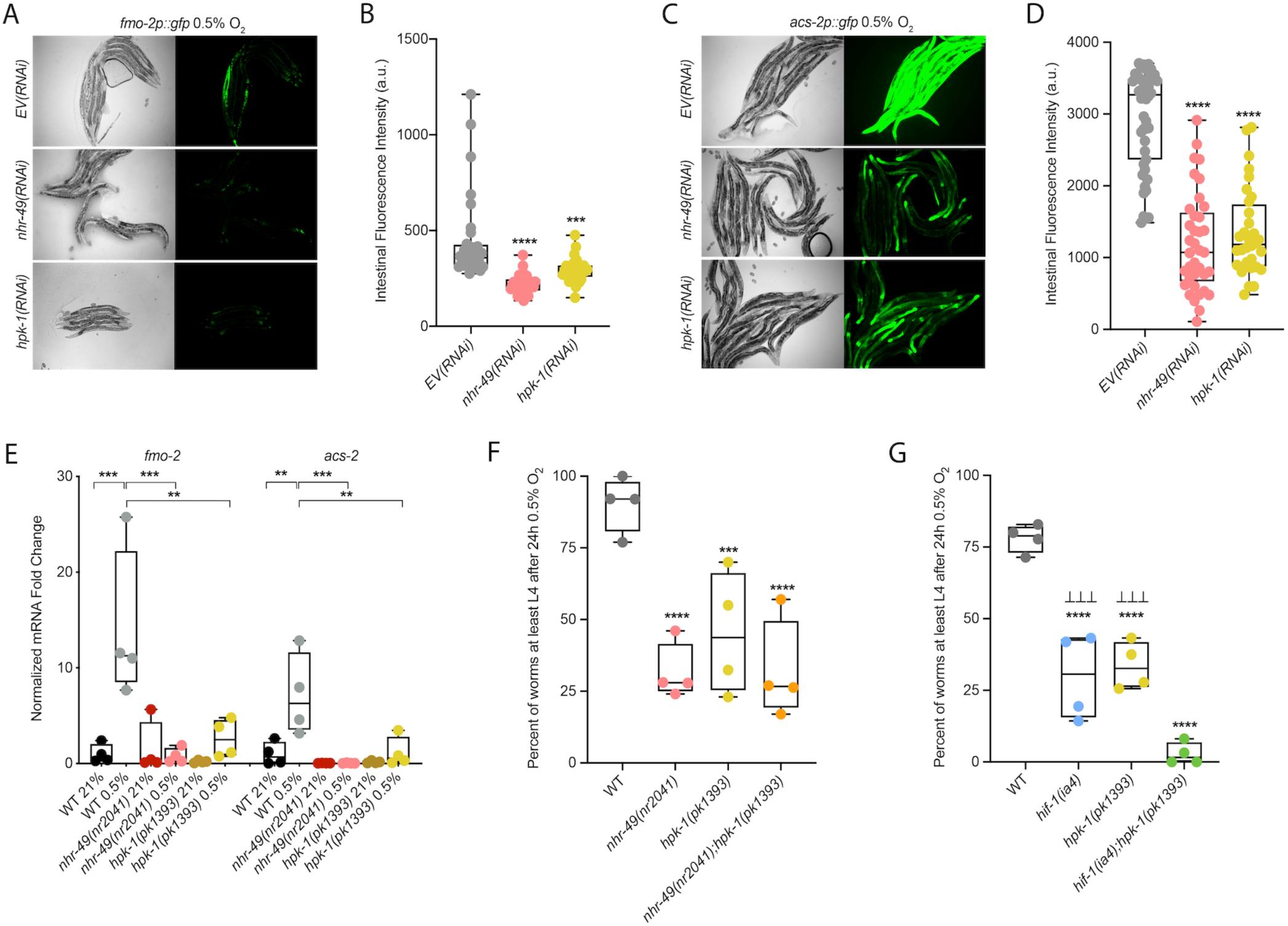
*hpk-1* is a positive regulator within the *nhr-49*-dependent hypoxia response pathway. **(A-D)** Representative micrographs and quantification of intestinal GFP levels in *Pfmo-2::gfp* (A, B) and *Pacs-2::gfp* (C, D) adult worms fed *EV*, *nhr-49*, *hif-1*, or *hpk-1* RNAi following 4 hr exposure to 0.5% O_2_ and 1 hr recovery in 21% O_2_ (3 or more repeats totalling >30 individual worms per strain). **,***,**** p <0.01, 0.001, 0.0001 vs. *EV(RNAi)* (ordinary one-way ANOVA corrected for multiple comparisons using the Tukey method). **(E)** The graph shows fold changes of mRNA levels in L4 wild-type, *nhr-49(nr2041)*, and *hpk-1(pk1393)* worms exposed to 0.5% O_2_ for 3 hr (n = 4). **,*** p < 0.01, 0.001 (ordinary one-way ANOVA corrected for multiple comparisons using the Tukey method). **(F)** The graph shows average population survival of wild-type, *nhr-49(nr2041)*, *hpk-1(pk1393)*, and *nhr-49(nr2041);hpk-1(pk1393)* worm embryos following 24 hr exposure to 0.5% O_2_, then allowed to recover at 21% O_2_ for 65 hr, and counted as ability to reach at least L4 stage (4 repeats totalling >100 individual worms per strain). ***,**** p<0.001, 0.0001 vs. wild-type worms. Comparison of single mutants to *nhr-49(nr2041);hpk-1(pk1393)* not significant (ordinary one-way ANOVA corrected for multiple comparisons using the Tukey method). **(G)** The graph shows average population survival of wild-type, *hif-1(ia4)*, *hpk-1(pk1393)*, and *hif-1(ia4);hpk-1(pk1393)* worm embryos following 24 hr exposure to 0.5% O_2_, then allowed to recover at 21% O_2_ for 65 hr, and counted as ability to reach at least L4 stage (four repeats totalling >100 individual worms per strain). **** p<0.0001 vs. wild-type worms, ⊥⊥⊥ p<0.001 vs. *hif-1(ia4);hpk-1(pk1393)* (ordinary one-way ANOVA corrected for multiple comparisons using the Tukey method). n.s. = not significant, WT = wild-type.

To determine if *hpk-1* is functionally required for worm survival in hypoxia, we assessed the ability of *hpk-1* mutant embryos to survive hypoxia. Similar to *nhr-49* mutant worms, only 45% of *hpk-1* mutant embryos developed to L4 (wild-type worms 92%; Figure 6F). We used epistasis analysis to test the hypothesis that *hpk-1* acts in the *nhr-49* pathway to coordinate a transcriptional response to hypoxia. We observed that the *nhr-49;hpk-1* double null mutant showed similar survival (26%) to each of the single null mutants, suggesting that these two genes act in the same hypoxia response pathway (Figure 6F). In contrast, the *hif-1;hpk-1* double null mutant was significantly impaired (<2%) compared to each of the single null mutants alone, consistent with the view that these two genes act in separate response pathways (Figure 6G). Each mutant showed normal development from embryo to L4 in normoxia, indicating that the phenotypes observed were specifically due to the requirement of these genes in hypoxia survival (Supplementary Figure 6C, D, Supplementary Table 1). Taken together, these experiments show that *hpk-1* is required for embryo survival in hypoxia, consistent with it playing a role as an activator of the *nhr-49*-dependent response pathway.

### NHR-49 is regulated post-transcriptionally in hypoxia in an *hpk-1*-dependent fashion

To test our hypothesis that HPK-1 activates NHR-49 in hypoxia, we examined whether NHR-49 is induced by hypoxia and whether *hpk-1* is involved in this regulation. NHR-49 and HPK-1 protein levels are increased in response to tert-butyl hydroperoxide and/or heat shock, respectively, but mRNA levels remain unchanged (Das et al., 2017; Goh et al., 2018). Similarly, we observed that *nhr-49* and *hpk-1* mRNA levels were not increased upon exposure to hypoxia (Figure 7A). Consistent with this, a transcriptional reporter of the *hpk-1* promoter fused to GFP (Das et al., 2017) was not induced following hypoxia exposure (Supplementary Figure 7A, B). These data show that transcription of neither *nhr-49* nor *hpk-1* are induced in hypoxia.

**Figure 7.**
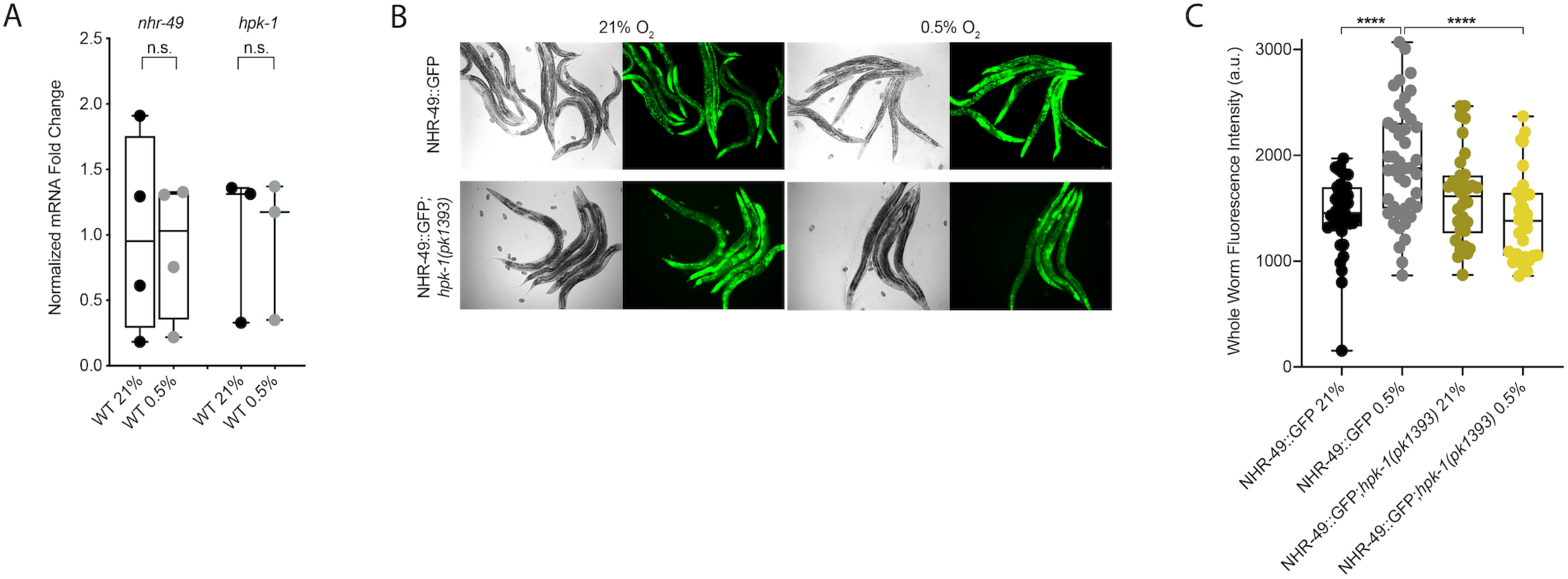
NHR-49 is induced in hypoxia in an *hpk-1*-dependent fashion. **(A)** The graph shows the average fold changes of mRNA levels (relative to unexposed wild-type wild-type) in L4 wild-type worms exposed to 0.5% O_2_ for 3 hr (n = 4; ordinary one-way ANOVA corrected for multiple comparisons using the Tukey method). **(B)** Representative micrographs show *Pnhr-49::nhr-49::gfp* and *Pnhr-49::nhr-49::gfp;hpk-1(pk1393)* adult worms following 4 hr exposure to 0.5% O_2_ and 1 hr recovery in 21% O_2_. **(C)** The graph shows the quantification of whole worm GFP levels in *Pnhr-49::nhr-49::gfp* and *Pnhr-49::nhr-49::gfp;hpk-1(pk1393)* worms following 4 hr exposure to 0.5% O_2_ and 1 hr recovery in 21% O_2_ (three repeats totalling >30 individual worms per strain). **** p <0.0001 (ordinary one-way ANOVA corrected for multiple comparisons using the Tukey method). n.s. = not significant, WT = wild-type.

We considered the possibility that NHR-49 may be regulated post-transcriptionally. To assess NHR-49 protein levels, we used the translational NHR-49::GFP reporter to measure the expression of the fusion protein in response to hypoxia. Indeed, the whole worm NHR-49::GFP signal was modestly, but significantly induced upon exposure to hypoxia (Figure 7B, C). Interestingly, while *hpk-1* null mutation had no effect on NHR-49::GFP levels in normoxia, it abrogated the induction of the NHR-49::GFP signal by hypoxia (Figure 7B, C). This suggests that NHR-49 is regulated post-translationally in response to hypoxia, and that *hpk-1* may be involved in this regulation. Taken together, these data show that *hpk-1* is a functionally important upstream positive regulator of the *nhr-49-*dependent hypoxia response.

## Discussion

Animals, tissues, and cells must be able to rapidly, flexibly, and reversibly adapt to a plethora of stresses. Past studies have identified many stress response factors, often termed master regulators. However, more recent studies indicate that stress response regulation requires the intricate interactions of multiple factors as part of networks that provide regulatory redundancy and flexibility. NHR-49 is a transcription factor that promotes longevity and development by regulating lipid metabolism and various stress responses (Chamoli et al., 2014; Dasgupta et al., 2020; Goh et al., 2018; Naim et al., 2020; Wani et al., 2020). Our data show that *nhr-49* coordinates a new aspect of the transcriptional response to hypoxia. This pathway operates in parallel to, and independent of, the canonical *hif-1* hypoxia response pathway. Besides *nhr-49*, it contains *nhr-67* and *hpk-1*. The former acts during normoxia to repress NHR-49; however, during hypoxia, an increase in NHR-49 protein levels in turn represses *nhr-67* levels, forming a feedback loop that may serve to reinforce NHR-49 activity. In contrast to *nhr-67*, the upstream kinase HPK-1 positively regulates the NHR-49 hypoxia response, as it is required to activate the NHR-49 regulated hypoxia response genes *fmo-2* and *acs-2* and to survive hypoxia. Downstream, NHR-49 induces autophagy genes, which are essential to promote hypoxia survival. Collectively, our experiments delineate a *hif-1*-independent hypoxia response pathway that contains distinct upstream and downstream components and is just as essential for hypoxia survival as is the *hif-1* pathway (Figure 8).

**Figure 8.**
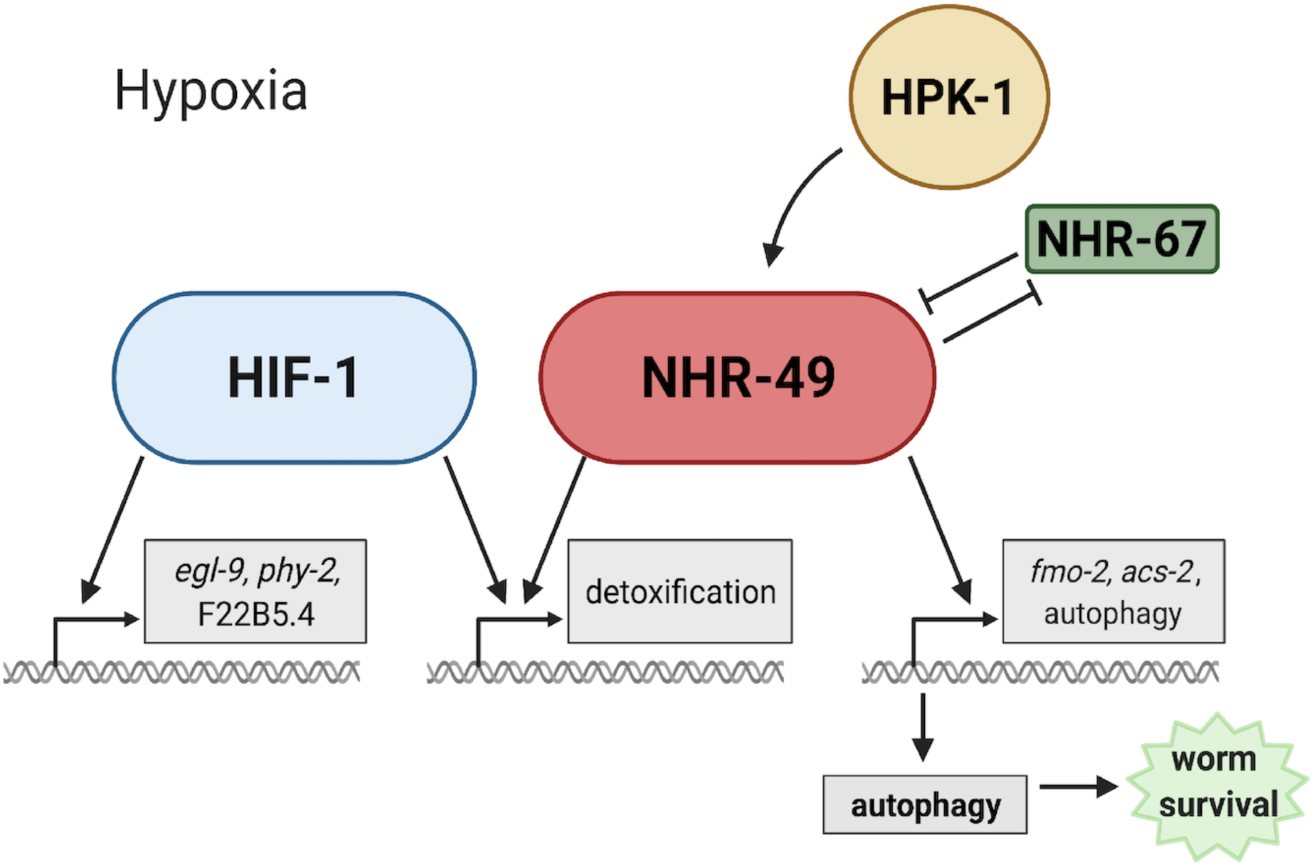
Model of the new NHR-49 hypoxia response pathway and its interaction with HIF-1 signaling. The proposed model of how NHR-49 regulates a new, *hif-1*-independent hypoxia response. During normoxia, the transcription factor NHR-67 negatively regulates NHR-49. However, during hypoxia, NHR-49 represses *nhr-67*, and the kinase HPK-1 positively regulates NHR-49. This allows NHR-49 to activate its downstream hypoxia response target genes, including *fmo-2*, *acs-2*, and autophagy genes, whose induction is required for worm survival to hypoxia. (Figure created with Biorender.com, Toronto, ON, Canada).

### NHR-49 controls a novel hypoxia response pathway that is parallel to canonical HIF signaling

*nhr-49* is required to induce *fmo-2* in various stresses and infection models (Chamoli et al., 2014; Dasgupta et al., 2020; Goh et al., 2018; Naim et al., 2020; Wani et al., 2020). Similarly, HIF-1 regulates *fmo-2* in several *C. elegans* longevity paradigms (Leiser et al., 2015), and *fmo-2* is induced in hypoxia, specifically 0.1% O_2_ exposure, in a *hif-1-*dependent manner (Leiser et al., 2015; Shen et al., 2005). This raised the possibility that *hif-1* also promoted *fmo-2* expression in hypoxia (0.5% O_2_) in L4 or older worms, and, more generally, that *nhr-49* might act through *hif-1* in the hypoxia response. However, several lines of evidence support a model whereby HIF-1 and NHR-49 are core components of parallel, independent signaling networks (Figure 8). First, *hif-1* and *nhr-49* interact genetically in hypoxia survival experiments, suggesting they work in separate genetic pathways (Figure 2A, B). Second, our transcriptome analysis identified sets of genes that are regulated exclusively by HIF-1 or NHR-49 (Figure 3A, B). Third, the kinase *hpk-1*, identified in a screen for new factors acting with NHR-49 in *fmo-2* induction, also shows synthetic genetic interaction with *hif-1*, but not with *nhr-49* (Figure 6F, G). Collectively these data show that *nhr-49* is a core part of a hypoxia response pathway that is independent of *hif-1* signalling. In support of our study, a recent publication (Vozdek et al., 2018) showed that *nhr-49* is required to induce the *hif-1*-independent hypoxia response gene *comt-5* both in 0.5% O_2_ and in a strain mutant for the kinase *hir-1*. In hypoxia, HIR-1 coordinates remodeling of the extracellular matrix independently of HIF-1 (Vozdek et al., 2018). Thus, although our RNA-seq results did not identify *comt-5* as a target of NHR-49 in hypoxia, this study supports the idea of a *hif-1*-independent hypoxia response pathway involving *nhr-49*.

### Homeodomain interacting protein kinases in hypoxia

Our efforts to map additional components of the NHR-49 hypoxia response pathway, especially factors acting in concert with NHR-49, revealed *hpk-1* (Figure 8). Homeodomain interacting protein kinases (HIPKs) are a family of nuclear serine/threonine kinase that can phosphorylate transcription factors (Rinaldo et al., 2007, 2008). The worm’s only HIPK orthologue, *hpk-1*, regulates development and the response to DNA damage, heat shock, and dietary restriction (Berber et al., 2013, 2016; Das et al., 2017; Rinaldo et al., 2007). Here, we show that *hpk-1* is an upstream regulator of the *nhr-49*-dependent hypoxia response pathway. Our data indicate that HPK-1 promotes the accumulation of NHR-49 protein in hypoxia, leading to induction of NHR-49-dependent hypoxia response genes. Interestingly, mammalian HIPK2 is degraded during periods of low oxygen via association with the E3 ubiquitin ligase SIAH2 (Calzado et al., 2009). This degradation of HIPK2 is necessary, as the protein normally represses the expression of HIF-1α by binding at its promoter in cell culture (Nardinocchi et al., 2009). In contrast, HIPK2 is induced in and required to protect cardiomyocytes from hypoxia/reoxygenation induced injury (Dang et al., 2020). This is consistent with our data and suggests that protecting cells from hypoxic injury may be a conserved role of HIPKs. Future experiments may reveal how HPK-1 is regulating NHR-49, perhaps examining direct phosphorylation and activation of the NHR-49 protein by HIPK-1.

### Paradoxical regulation of the β-oxidation gene *acs-2* by hypoxia

Mitochondria consume cellular oxygen to produce energy and thus must adapt to limited oxygen availability. In particular, mitochondrial β-oxidation, the consumption of oxygen to catabolize fatty acids for energy production, is repressed in hypoxia in favour of anaerobic respiration. For example, the heart and skeletal muscle of mice and rats show decreased expression of key β-oxidation enzymes in acute hypoxia (Kennedy et al., 2001; Morash et al., 2013). In *C. elegans*, the acyl-CoA synthetase *acs-2* is part of the mitochondrial β-oxidation pathway, where it functions in the first step to activate fatty acids. NHR-49 activates *acs-2* expression during starvation, when β-oxidation is induced (M. R. Van Gilst et al., 2005). Considering this, *acs-2* expression would be expected to be downregulated in hypoxia due to reduced β-oxidation. Paradoxically, however, we found that *acs-2* is strongly induced in hypoxia and that this regulation depends on *nhr-49* (Figure 3C, Supplementary Figure 3A-C). Examination of other fatty acid β-oxidation enzymes in our RNA-seq data showed that *acs-2* is the only enzyme induced. This suggests that, during hypoxia, ACS-2 is not feeding its product fatty acyl-CoA into the β-oxidation cycle, but perhaps produces fatty acyl-CoA for anabolic functions needed for survival to or recovery from low oxygen, such as phospholipid or triglyceride synthesis (reviewed in Tang et al., 2018). Similar functions have been observed in human macrophages, which, during hypoxia, decrease β-oxidation but increase triglyceride synthesis (Boström et al., 2006).

In line with the repression of β-oxidation in hypoxia (Boström et al., 2006; Kennedy et al., 2001; Morash et al., 2013), there is evidence supporting a HIF-dependent down-regulation of the mammalian NHR-49 homolog PPARα, which promotes β-oxidation (Atherton et al., 2008). For example, in human hepatocytes and in mouse liver sections, HIF-2α accumulation in hypoxia directly suppresses PPARα expression (J. Chen et al., 2019). Additionally, HIF-1α suppresses PPARα protein and mRNA levels during hypoxia in intestinal epithelial cells, and the *PPARA* promoter contains a HIF-1α DNA binding consensus motif, suggesting direct control of *PPARA* by HIF transcription factors (Narravula & Colgan, 2001).

Some evidence suggests alternative actions of PPARα. Knockdown of PPARα attenuates the ability of Phd1 (the homolog of *C. elegans egl-9*) knockout myofibers to successfully tolerate hypoxia (Aragonés et al., 2008), suggesting that PPARα is an important regulator of the hypoxia response downstream of Phd1. Along these lines, PPARα protein levels increase in the muscle of Phd1 knockout mice (Aragonés et al., 2008) and following hypoxic exposure in mouse hearts (Morash et al., 2013). Similarly, we show that NHR-49 protein levels increase in response to hypoxia (Figure 7B, C), and that NHR-49 is a vital regulator of a *hif-1*-independent hypoxia response. Together, these data suggest that, similar to evidence from studies in mammalian systems, NHR-49 levels are increased and required in hypoxia, and may be regulating *acs-*2 for functions other than fatty acid β-oxidation.

### NHR-49 promotes autophagy activation to achieve hypoxia survival

Damaged cellular components can be cleared via autophagy, a key process regulated by *nhr-49* in hypoxia (Figure 3D, F). In mammals, PPARα activates autophagy in response to various stresses, including in neurons to clear Aβ in Alzheimer’s disease (Luo et al., 2020), and in the liver during inflammation (Jiao et al., 2014) and starvation (J. M. Lee et al., 2014). Proper regulation of autophagy is also a requirement in hypoxic conditions. Knockdown or genetic mutation of various *C. elegans* autophagy genes showed that they are required for worm survival when worms experience anoxia and elevated temperatures combined (Samokhvalov et al., 2008). Similarly, Zhang *et al*. found that mitochondrial autophagy (mitophagy) is induced by hypoxia in mouse embryo fibroblasts. This process requires expression of BNIP3 (Bcl-2/E1B 19 kDa-interacting protein 3), an autophagy inducer, which is induced in a HIF-1-dependent manner (H. Zhang et al., 2008). In agreement with this, our RNA-seq data showed a 3.8-fold induction of the *C. elegans* BNIP3 homolog *dct-1* in hypoxia; however, this induction was dependent on neither *nhr-49* nor *hif-1*. The above study also found that the autophagy genes Beclin-1 and Atg5 are induced and required for cell survival in hypoxia (H. Zhang et al., 2008). Here, we show that the *C. elegans* ortholog of Beclin-1, *bec-1*, and the worm *atg-7* and *atg-10* genes, which are involved in the completion of the autophagosome along with *atg-5/Atg5,* are required for worm embryo survival to hypoxia in an *nhr-49*-dependent manner (Figure 3F). Interestingly, *hpk-1* regulates autophagy in response to dietary restriction, as it is necessary to induce autophagosome formation and autophagy gene expression (Das et al., 2017). *hpk-1* may thus aid *nhr-49* in the regulation of autophagy during hypoxia as well.

### Cell non-autonomous functions of NHR-49 in hypoxia

Cell non-autonomous regulation occurs in many pathways in *C. elegans*. For example, HIF-1 acts in neurons to induce *fmo-2* expression in the intestine to promote longevity (Leiser et al., 2015). NHR-49 is expressed in the intestine, neurons, muscle, and hypodermis (Ratnappan et al., 2014). Re-expression of *nhr-*49 in any one of these tissues is sufficient to enhance worm survival upon infection with the pathogens *S. aureus* (Wani et al., 2020) and to promote longevity in germline-less animals (Naim et al., 2020), but NHR-49 acts only in neurons to promote survival to *P. aeruginosa* (Naim et al., 2020). We thus aimed to identify the key tissue wherein NHR-49 promotes hypoxia survival. Surprisingly, we found that *nhr-49* expression in any of the intestine, neurons, or hypodermis is sufficient for whole animal survival to hypoxia (Figure 4B), suggesting that NHR-49 can act in a cell non-autonomous fashion to execute its effects. Possibly, a signaling molecule whose synthesis is promoted by NHR-49 activity in any tissue promotes organismal hypoxia adaptation.

In sum, we show here that NHR-49 regulates a novel hypoxia response pathway that is independent of HIF-1 and controls an important transcriptional response for worm survival to hypoxia. If the mammalian NHR-49 homologs PPARα and HNF4 play similar roles in the cellular response to hypoxia, our discovery could lead to the identification and development of new targets for drugs and therapies for diseases exhibiting hypoxic conditions.

## Materials and Methods

### Nematode strains and growth conditions

We cultured *C. elegans* strains using standard techniques on nematode growth media (NGM) plates. To avoid background effects, each mutant was crossed into our lab N2 strain; original mutants were backcrossed to N2 at least six times. *E. coli* OP50 was the food source in all experiments except for RNAi experiments, where we used *E. coli* HT115. All experiments were carried out at 20°C. Worm strains used in this study are listed in Supplementary Table 6. For synchronized worm growths, we isolated embryos by standard sodium hypochlorite treatment. Isolated embryos were allowed to hatch overnight on unseeded NGM plates until the population reached a synchronized halted development at L1 stage via short-term fasting (12–24 hr). Synchronized L1 stage larvae were then transferred to OP50 seeded plates and grown to the desired stage.

### Feeding RNA interference

RNAi was performed on NGM plates supplemented with 25 μg/ml carbenicillin (BioBasic CDJ469), 1 mM IPTG (Santa Cruz CAS 367-93-1), and 12.5 μg/ml tetracycline (BioBasic TB0504 (NGM-RNAi plates), and seeded with appropriate HT115 RNAi bacteria. The RNAi clones were from the Ahringer library (Source BioScience) and were sequenced prior to use.

### RNA isolation and qRT-PCR analysis

Synchronized L1 worms were allowed to grow on OP50 plates for 48 hr to L4 stage, then either kept in 21% O_2_ or transferred to 0.5% O_2_ for 3 hr. RNA isolation was performed as previously described (Goh et al., 2014). 2 μg total RNA was used to generate cDNA with Superscript II reverse transcriptase (Invitrogen 18064-014), random primers (Invitrogen 48190-011), dNTPs (Fermentas R0186), and RNAseOUT (Invitrogen 10777-019). Quantitative PCR was performed in 10 μl reactions using Fast SYBR Master Mix (Life Technologies 4385612), 1:10 diluted cDNA, and 5 μM primer, and an analyzed with an Applied Biosystems StepOnePlus machine. We analyzed the data with the ΔΔCt method. For each sample, we calculated normalization factors by averaging the (sample expression)/(average reference expression) ratios of three normalization genes, *act-1*, *tba-1*, and *ubc-2*. The reference sample was *EV(RNAi)*, wild-type, or 21% O_2_, as appropriate. We used one-way ANOVA to calculate statistical significance of gene expression changes and corrected for multiple comparisons using the Tukey method. Primers were tested on serial cDNA dilutions and analyzed for PCR efficiency prior to use. All data originate from three or more independent biological repeats, and each PCR reaction was conducted in technical triplicate. Sequences of qRT-PCR primers are listed in Supplementary Table 7.

### Analysis of fluorescent reporter lines via DIC and fluorescence microscopy

To analyze fluorescence in reporter lines, egg lays were performed on NGM plates seeded with OP50 or RNAi plates seeded with the appropriate HT115 RNAi culture. Worms were allowed to grow to adulthood. Plates were then kept in 21% O_2_ or transferred to 0.5% O_2_ for 4 hr and allowed to recover for 1 hr in normoxia before imaging. Worms were collected into M9 buffer containing 0.06% levamisole (Sigma L9756) for immobilization on 2% (w/v) agarose pads for microscopy. We captured images on a CoolSnap HQ camera (Photometrics) attached to a Zeiss Axioplan 2 compound microscope, followed by MetaMorph Imaging Software with Autoquant 3D digital deconvolution. All images for the same experiment were captured at the same exposure time. Images were analyzed using ImageJ software (https://imagej.nih.gov/ij/download.html), with fluorescence calculated by taking the difference of the background fluorescence from the mean intestinal or whole worm fluorescence. For experiments imaging the *Pfmo-2::gfp* and *Pacs-2::gfp* reporters, intestinal fluorescence was measured. For experiments imaging the *Phpk-1::gfp*, *Pnhr-49::nhr-49::gfp*, or the *Phpk-1::hpk-1::gfp* reporters, whole worm fluorescence was measured. For each experiment, at least three independent trials were performed with a minimum of 30 worms per condition.

### NHR-49 transgenic strains

To construct the *Pnhr-49::nhr-49::gfp* containing plasmid, a 6.6 kb genomic fragment of the *nhr-49* gene (including a 4.4 kb coding region covering all *nhr-49* transcripts and a 2.2 kb promoter region) was cloned into the GFP expression vector pPD95.77 (Addgene #1495), as reported previously (Ratnappan et al., 2014). For generating tissue-specific constructs, the *nhr-49* promoter was replaced with tissue-specific promoters using *Sbf*I and *Sal*I restriction enzymes to create plasmids for expressing NHR-49 in the muscle (*Pmyo-3::nhr-49::gfp*), intestine (*Pgly-19::nhr-49::gfp*), hypodermis (*Pcol-12::nhr-49::gfp*), and neurons (*Prgef-1::nhr-49::gfp*). 100 ng/μl of each plasmid was injected, along with pharyngeal muscle-specific *Pmyo-2::mCherry* as a co-injection marker (25 ng/μl) into the *nhr-49(nr2041)* mutant strain using standard methods (Mello & Fire, 1995). Strains were maintained by picking animals that were positive for both GFP and mCherry.

### Hypoxia sensitivity assays

Hypoxic conditions were maintained using continuous flow chambers, as previously described (Fawcett et al., 2012). Compressed gas tanks (5000 ppm O_2_ balanced with N_2_) were certified as standard to within 2% of indicated concentration from Praxair Canada (Delta, BC). Oxygen flow was regulated using Aalborg rotameters (Aalborg Instruments and Controls, Inc., Orangeburg, NY, USA). Hypoxic chambers (and room air controls) were maintained in a 20°C incubator for the duration of the experiments.

For embryo survival assays, gravid first-day adult worms (picked as L4 the previous day) were allowed to lay eggs for 1-4 hr on plates seeded with 15 uL OP50 or appropriate HT115 RNAi bacteria the previous day. Adults were removed, and eggs were exposed to 0.5% O_2_ for 24 hr or 48 hr. Animals were scored for developmental success (reached at least L4 stage) after being placed back into room air for 65 hr (following 24 hr exposure) or 42 hr (following 48 hr exposure). For RNAi survival assays, worms were grown for one generation from egg to adult on the appropriate HT115 RNAi bacteria before their progeny was used for the egg lay.

For larval development assays, gravid adult worms (picked as L4 the previous day) were allowed to lay eggs for 2 hr and kept at 20°C for 13-17 hr to allow hatching (egg lays for *nhr-49(nr2041)* strains with embryonic developmental delays were performed 2 hr earlier to ensure synchronization with wild-type worms). Freshly hatched L1 worms were transferred to plates seeded with 15μL OP50 the previous day, and exposed to 0.5% O_2_ for 48 hr. Animals were placed back into room air and immediately scored for stage.

For all normoxia (21% O_2_) comparison experiments, methods were as described above except plates were kept in room air for the duration (instead of being exposed to 0.5% O_2_).

### Hydrogen sulfide sensitivity assay

Construction of H_2_S chambers was as previously described (Fawcett et al., 2012; Miller & Roth, 2007). In short, 5000 ppm H_2_S (balanced with N_2_) was diluted with room air to a final concentration of 50 ppm and monitored with a custom H_2_S detector, as described (Miller & Roth, 2007). Compressed gas mixtures were obtained from Airgas (Seattle, WA) and certified as standard to within 2% of the indicated concentration. Survival assays were performed with 20 L4 animals picked onto OP50 seeded plates. Plates were exposed to 50 ppm H_2_S for 24 hr in a 20°C incubator, then returned to room air to score viability. Animals were scored 30 min after removal from H_2_S, and plates with dead animals were re-examined after several hours to ensure animals had not reanimated.

### RNA sequencing

Synchronized L1 wild-type, *nhr-49(nr2041)*, and *hif-1(ia4)* worms were allowed to grow on OP50 plates to L4 stage, then either kept in 21% O_2_ or transferred to 0.5% O_2_ for 3 hr. RNA was isolated from whole worms as described above. RNA integrity and quality were ascertained on a BioAnalzyer. Construction of strand-specific mRNA sequencing libraries and sequencing (75bp PET) on an Illumina HiSeq 2500 machine was done at the Sequencing Services facility of the Genome Sciences Centre, BC Cancer Agency, Vancouver BC, Canada (https://www.bcgsc.ca/services/sequencing-services). The raw FASTQ reads obtained from the facility were trimmed using Trimmomatic version 0.36 (Bolger et al., 2014) with parameters LEADING:3 TRAILING:3 SLIDINGWINDOW:4:15 MINLEN:36. Next, the trimmed reads were aligned to the NCBI reference genome WBcel235 WS277 (https://www.ncbi.nlm.nih.gov/assembly/GCF_000002985.6/) using Salmon version 0.9.1 (Patro et al., 2017) with parameters -l A -p 8 --gcBias. Then, transcript-level read counts were imported into R and summed into gene-level read counts using tximport (Soneson et al., 2015). Genes not expressed at a level greater than 1 count per million (CPM) reads in at least three of the samples were excluded from further analysis. The gene-level read counts were normalized using the trimmed mean of M-values (TMM) in edgeR (Robinson et al., 2010) to adjust samples for differences in library size. Differential expression analysis was performed using the quasi-likelihood F-test with the generalized linear model (GLM) approach in edgeR (Robinson et al., 2010). Differentially expressed genes (DEGs) were defined as those with at least two-fold difference between two individual groups at a false discovery rate (FDR) < 0.05. RNA-seq data have been deposited at NCBI Gene Expression Omnibus (https://www.ncbi.nlm.nih.gov/geo/) under the record GSE166788.

Functional enrichment analysis and visualization were performed using the Overrepresentation Analysis (ORA) module with the default parameters in easyGSEA in the eVITTA suite (https://tau.cmmt.ubc.ca/eVITTA/; Cheng X., Yan J., *et al*., in preparation). easyVizR in the eVITTA suite was used to visualize the overlaps and disjoints in the DEGs (input December 14, 2020).

## Acknowledgements

We thank the Taubert, Miller, and Ghazi labs for comments on the manuscript. Some strains were provided by the CGC, which is funded by NIH Office of Research Infrastructure Programs (P40 OD010440). Grant support was from The Canadian Institutes of Health Research (CIHR; PJT-153199 to ST), the Natural Sciences and Engineering Research Council of Canada (NSERC; RGPIN-2018-05133 to ST), the Cancer Research Society (CRS; to ST), and the National Institutes of Health (NIH; R01AG051659 to AG, R01AG044378 to DM). KRSD was supported by NSERC CGS-M, NSERC CGS-D, and BCCHR scholarships, and ST by a Canada Research Chair.

## Supplementary Figures

**Supplementary Figure 1.**
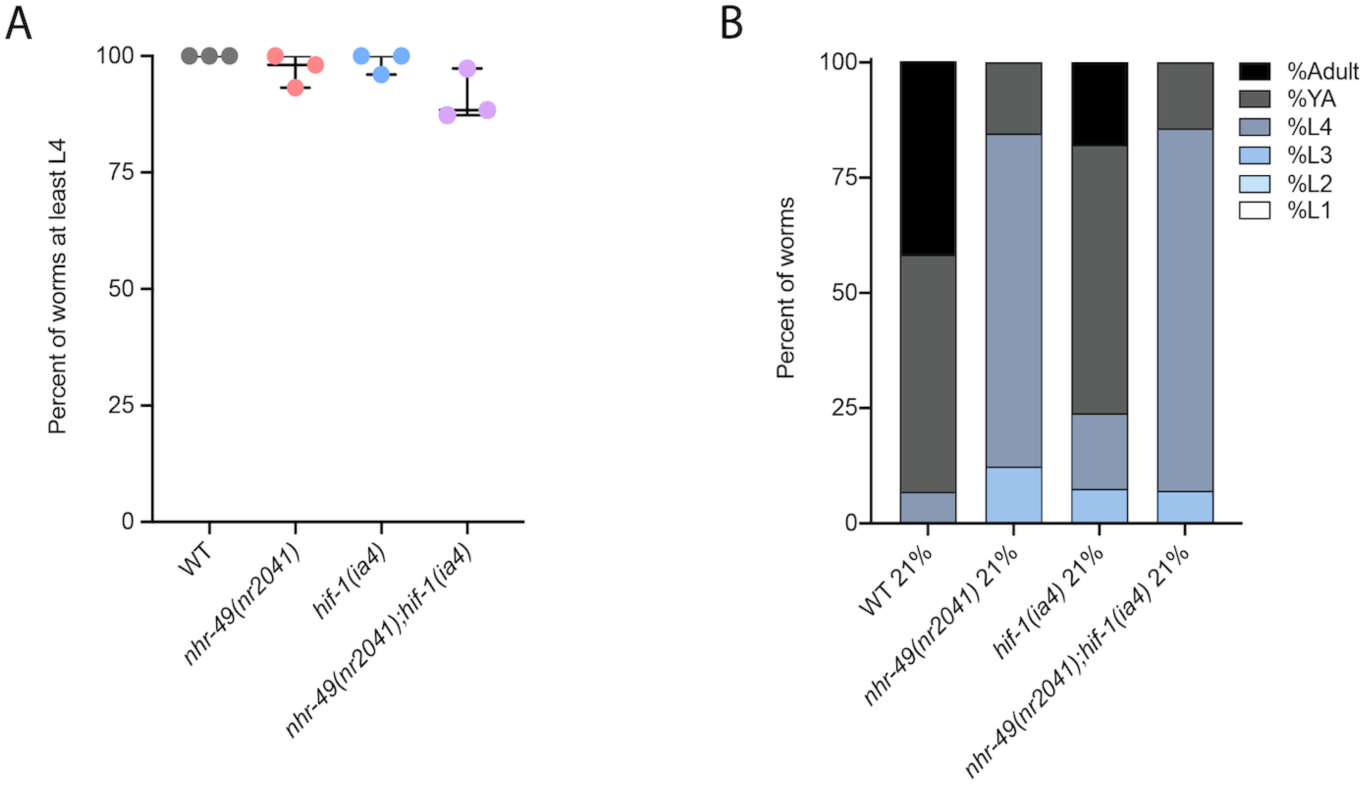
*nhr-49* and *hif-1* mutants do not display developmental defects in normoxia. **(A)** The graph shows the average developmental success of wild-type, *nhr-49(nr2041)*, *hif-1(ia4)*, and *nhr-49(nr2041);hif-1(ia4)* worm embryos kept in 21% O_2_ for 65 hr, and counted as ability to reach at least L4 stage (three repeats totalling >100 individual worms per strain). All comparisons not significant (ordinary one-way ANOVA corrected for multiple comparisons using the Tukey method). **(B)** The graph shows the average developmental success of wild-type, *nhr-49(nr2041)*, *hif-1(ia4)*, and *nhr-49(nr2041);hif-1(ia4)* larval worms kept in 21% O_2_ for 48 hr from L1 stage (four repeats totalling >60 individual worms per strain). All comparisons not significant (ordinary one-way ANOVA corrected for multiple comparisons using the Tukey method). WT = wild-type.

**Supplementary Figure 2.**
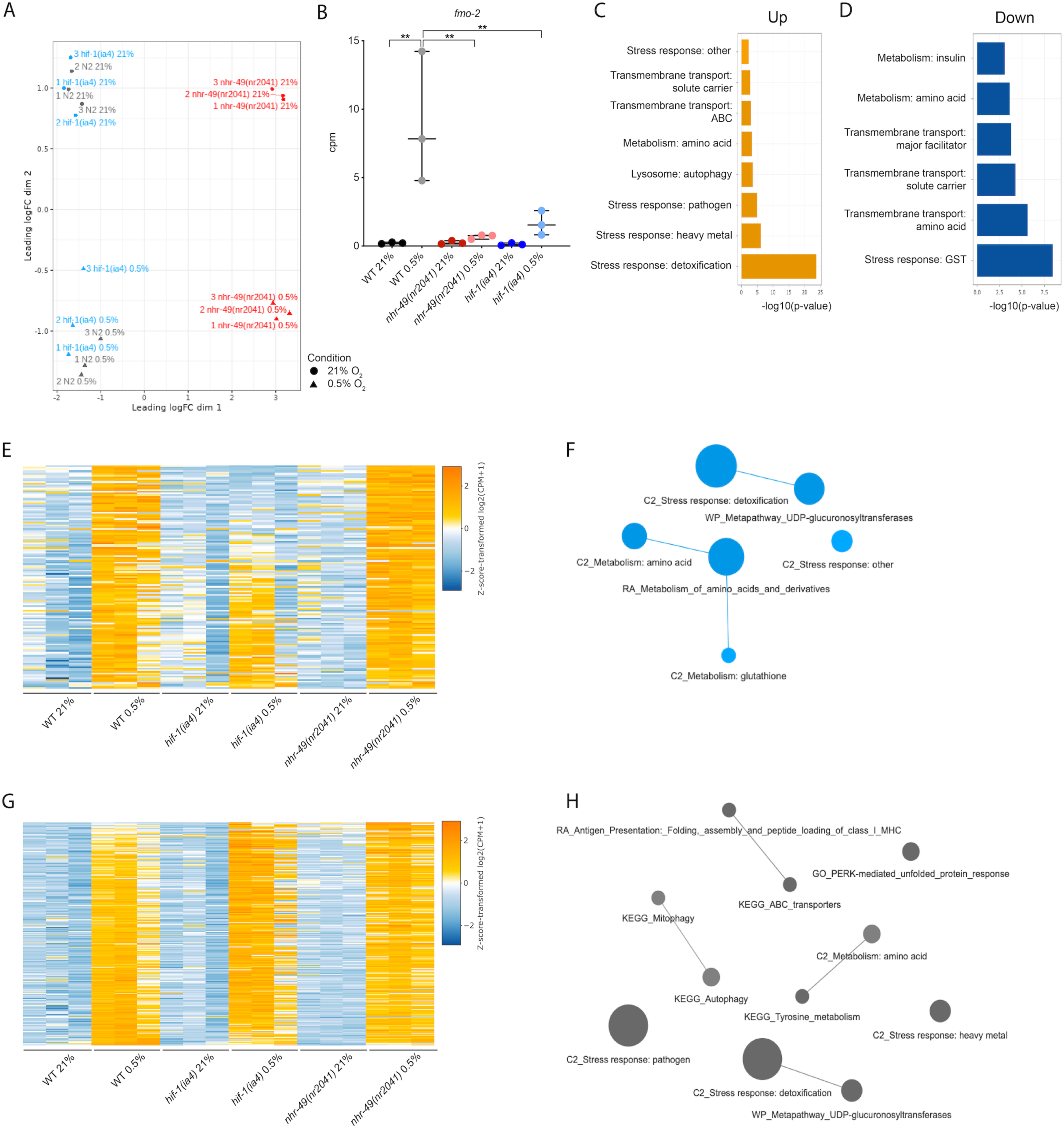
RNA-seq reveals discrete hypoxia responsive transcriptional programs. **(A)** The figure shows a Multi-Dimensional Scaling (MDS) plot of the distances between gene expression profiles. Distances on the MDS plot correspond to the root-mean-square average of the largest 200 log2-fold-changes between each pair of samples. **(B)** The graph shows average transcript levels in counts per million (CPM) of *fmo-2* mRNA in L4 wild-type, *nhr-49(nr2041)*, and *hif-1(ia4)* worms exposed to 0.5% O_2_ for 3 hr or kept at 21% O_2_ (n = 3). ** p <0.01 (ordinary one-way ANOVA corrected for multiple comparisons using the Tukey method). **(C, D)** Enriched WormCat (Category 2) categories among genes that are significantly up-regulated over two-fold (C) or down-regulated over two-fold (D) in wild-type worms in 21% O_2_ vs. 0.5% O_2_ are plotted by -log10 p-value. **(E, G)** Heatmaps of the expression levels of the (E) 139 genes from three repeats which are significantly induced over two-fold in 21% O_2_ vs. 0.5% O_2_ in wild-type and *nhr-49(nr2041)* worms, but not in *hif-1(ia4)*, and (G) the 264 genes from three repeats which are significantly induced over two-fold in 21% O_2_ vs. 0.5% O_2_ in wild-type, *hif-1(ia4)*, and *nhr-49(nr2041)* worms. Genes along the y-axis are colored in each repeat based on their z-scores of the log2-transformed Counts Per Million (CPM) plus 1. **(F, H)** Network views of the enriched functional categories among the 139 genes which are significantly induced over two-fold in 21% O_2_ vs. 0.5% O_2_ in wild-type and *nhr-49(nr2041)* worms, but not in *hif-1(ia4)* (F), and the 264 genes which are significantly induced over two-fold in 21% O_2_ vs. 0.5% O_2_ in wild-type, *hif-1(ia4)*, and *nhr-49(nr2041)* worms (H). Edge represents significant gene overlap as defined by a Jaccard Coefficient larger than or equal to 25%. Dot size reflects number of genes in each functional category; colour intensity reflects statistical significance (−log10 p-value). WT = wild-type.

**Supplementary Figure 3.**
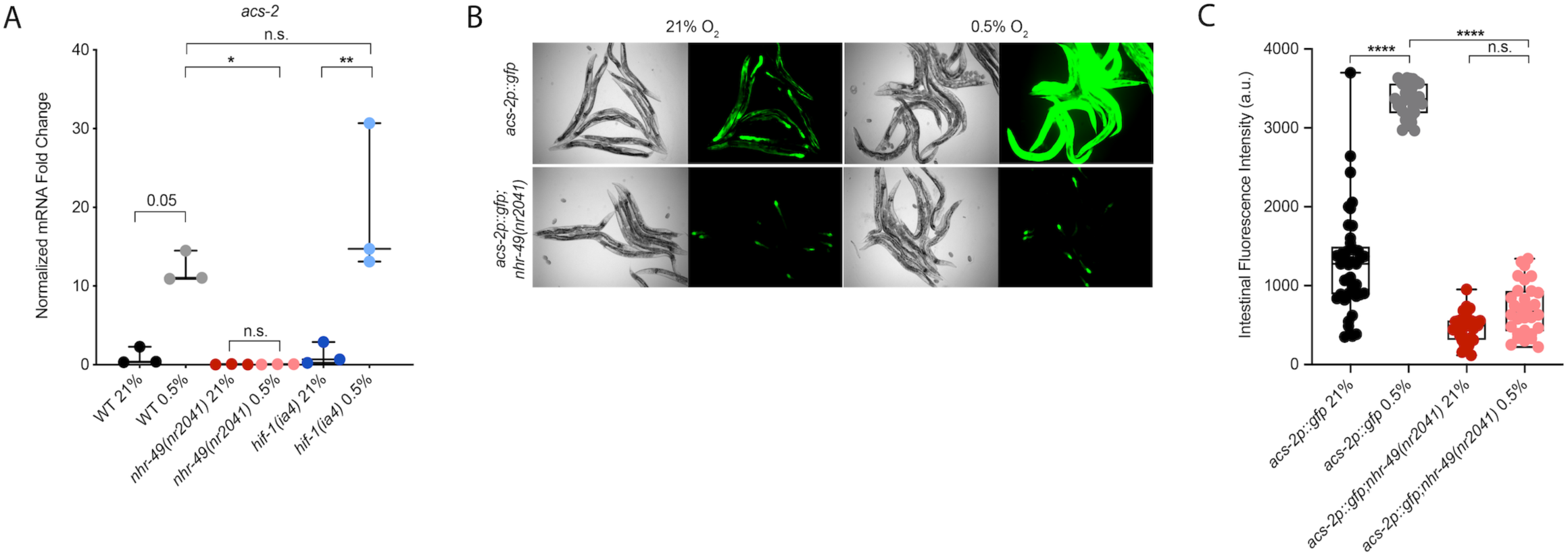
*nhr-49* regulates *acs-2* induction following exposure to hypoxia. **(A)** The graph shows average fold changes of mRNA levels (relative to unexposed wild-type) in L4 wild-type, *nhr-49(nr2041)*, and *hif-1(ia4)* worms exposed to 0.5% O_2_ for 3 hr (n = 3). *,** p < 0.05, (ordinary one-way ANOVA corrected for multiple comparisons using the Tukey method). **(B)** Representative micrographs show *Pacs-2::gfp* and *Pacs-2::gfp;nhr-49(nr2041)* adult worms following 4 hr exposure to 0.5% O_2_ and 1 hr recovery in 21% O_2_. **(C)** The graph shows the quantification of intestinal GFP levels in *Pacs-2::gfp* and *Pacs-2::gfp;nhr-49(nr2041)* worms following 4 hr exposure to 0.5% O_2_ and 1 hr recovery in 21% O_2_ (three repeats totalling >30 individual worms per strain). **** p <0.0001 (ordinary one-way ANOVA corrected for multiple comparisons using the Tukey method). n.s. = not significant, WT = wild-type.

**Supplementary Figure 4.**
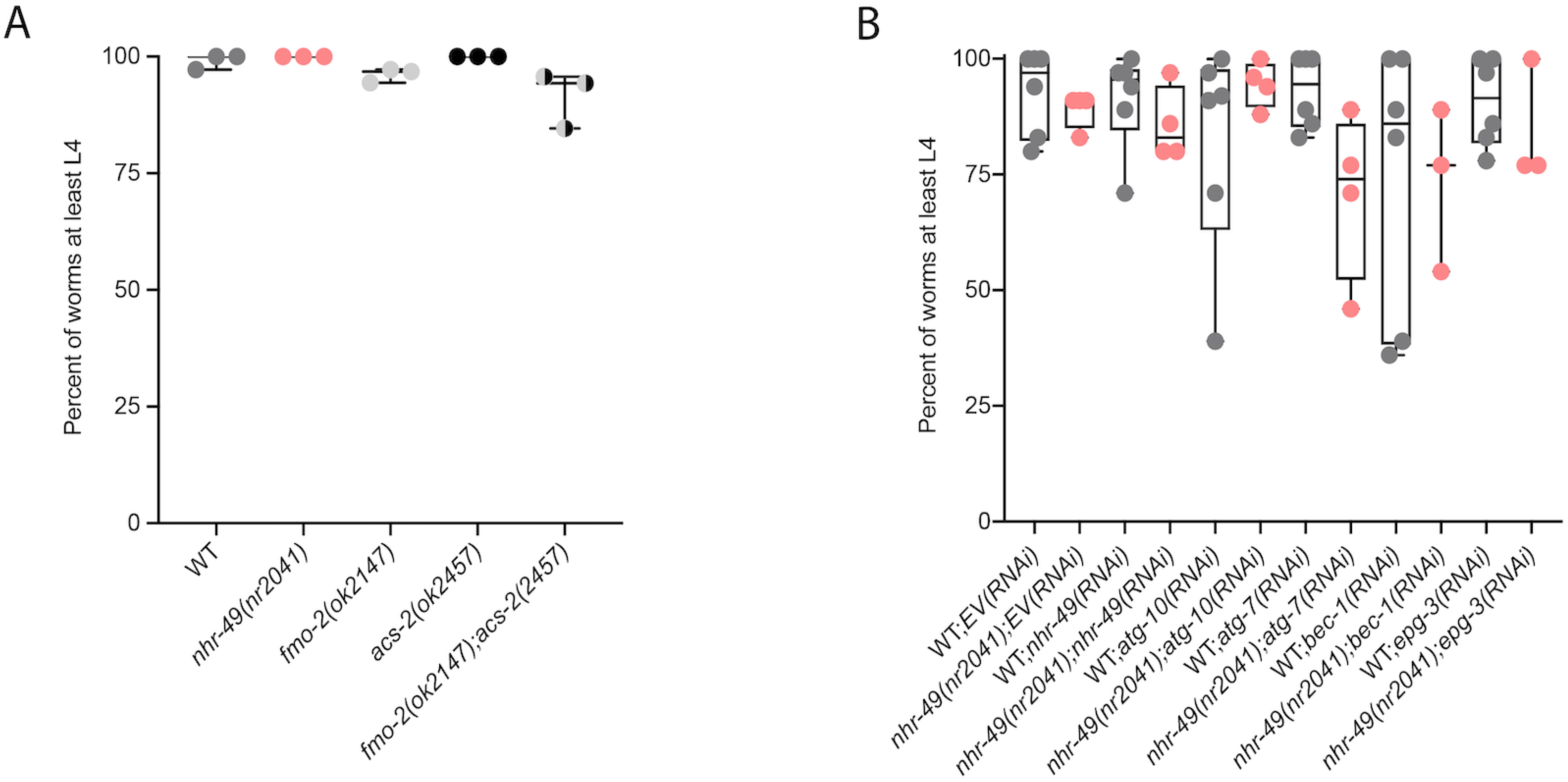
Mutants of downstream transcriptional targets of *nhr-49* in hypoxia do not display functional defects in normoxia. **(A)** The graph shows the average population survival of wild-type, *nhr-49(nr2041)*, *fmo-2(ok2147)*, *acs-2(ok2457)*, and *fmo-2(ok2147);acs-2(ok2457)* worm embryos kept in 21% O_2_ for 65 hr, and counted as ability to reach at least L4 stage (three repeats totalling >100 individual worms per strain). All comparisons not significant (ordinary one-way ANOVA corrected for multiple comparisons using the Tukey method). **(B)** The graph shows the average population survival of second generation wild-type and *nhr-49(nr2041)* worm embryos fed EV, *nhr-49*, *atg-10*, *atg-7*, *bec-1*, or *epg-3* RNAi kept in 21% O_2_ for 65 hr, and counted as ability to reach at least L4 stage (three repeats totalling >100 individual worms per strain). All comparisons not significant (ordinary one-way ANOVA corrected for multiple comparisons using the Tukey method). WT = wild-type.

**Supplementary Figure 5.**
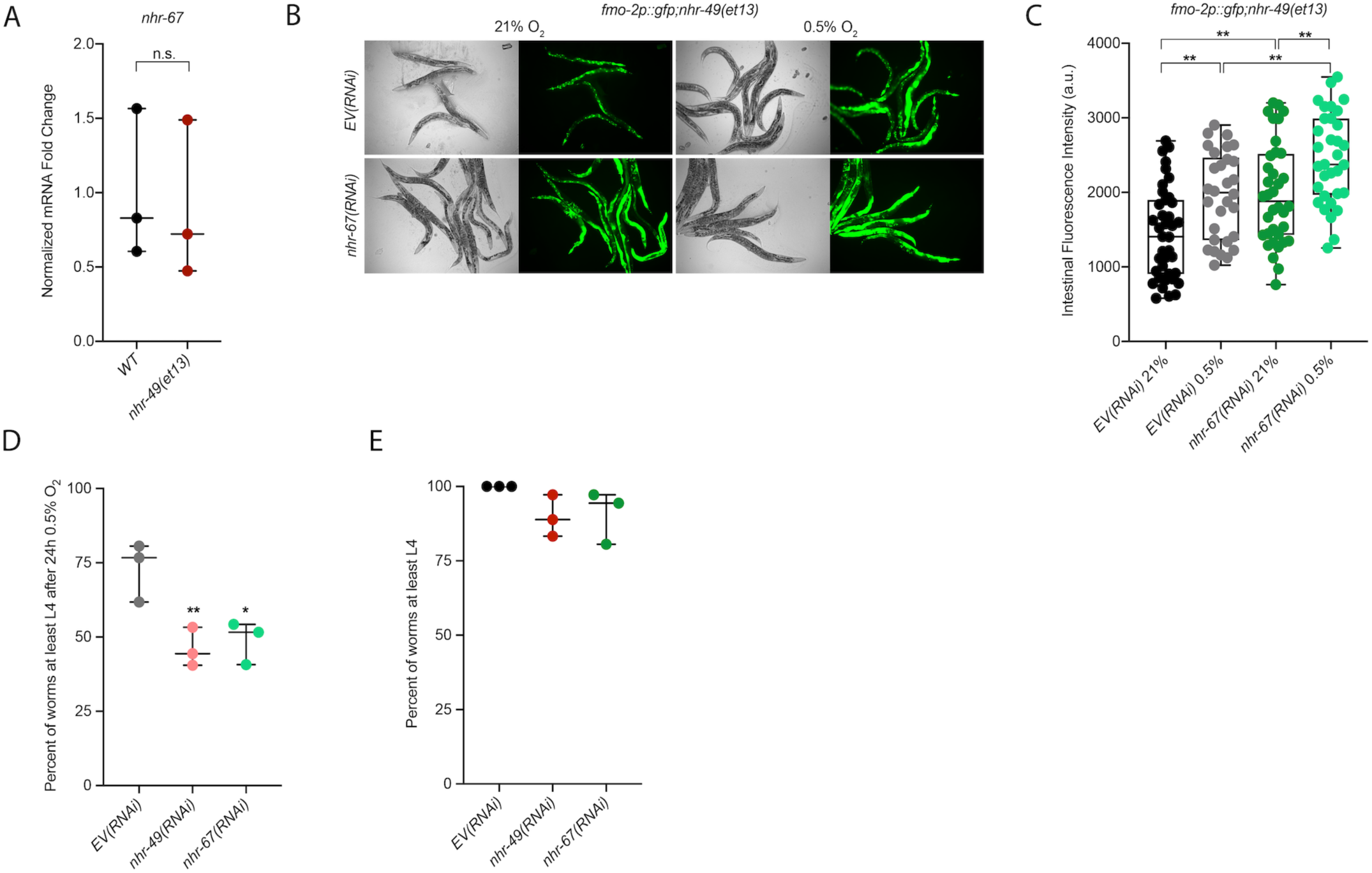
*nhr-67* is functionally required for survival in hypoxia. **(A)** The graph shows average fold changes of mRNA levels (relative to wild-type) in L4 wild-type and *nhr-49(et13)* worms (n = 3; ordinary one-way ANOVA corrected for multiple comparisons using the Tukey method). **(B-C)** Representative micrographs (B) and quantification (C) of intestinal GFP levels in *Pfmo-2::gfp;nhr-49(et13)* adult worms fed EV or *nhr-67* RNAi kept in 21% O_2_ (three repeats totalling >30 individual worms per strain). ** p <0.01 (ordinary one-way ANOVA corrected for multiple comparisons using the Tukey method). **(D)** The graph shows average population survival of second generation wild-type worm embryos fed EV, *nhr-49*, or *nhr-67* RNAi following 24 hr exposure to 0.5% O_2_, then allowed to recover at 21% O_2_ for 65 hr, and counted as ability to reach at least L4 stage (four repeats totalling >100 individual worms per strain). *, ** p<0.05,0.01 vs. *EV(RNAi)* worms (ordinary one-way ANOVA corrected for multiple comparisons using the Tukey method). **(E)** The graph shows the average population survival of second generation wild-type worm embryos fed EV, *nhr-49*, or *nhr-67* RNAi kept in 21% O_2_ for 65 hr, and counted as ability to reach at least L4 stage (four repeats totalling >100 individual worms per strain). All comparisons not significant (ordinary one-way ANOVA corrected for multiple comparisons using the Tukey method). n.s. = not significant, WT = wild-type.

**Supplementary Figure 6.**
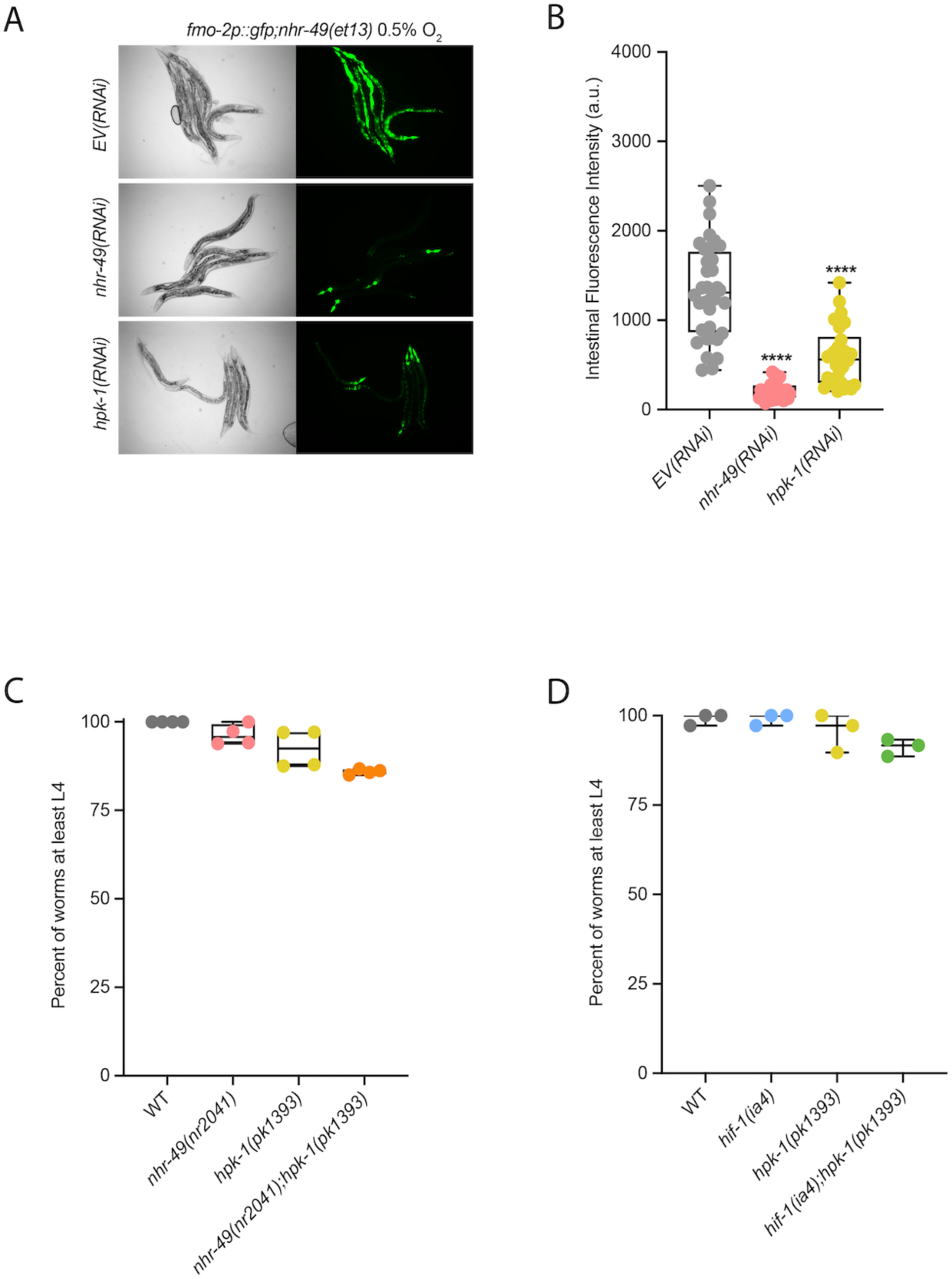
*hpk-1* mutants do not display functional defects in normoxia. **(A, B)** Representative micrographs (A) and quantification (B) of intestinal GFP levels in *Pfmo-2::gfp;nhr-49(et13)* adult worms fed EV, *nhr-49*, *hif-1*, or *hpk-1* RNAi kept in 21% O_2_ (three or more repeats totalling >30 individual worms per strain). **** p <0.0001 vs. *EV(RNAi)* (ordinary one-way ANOVA corrected for multiple comparisons using the Tukey method). **(C)** The graph shows average population survival of wild-type, *nhr-49(nr2041)*, *hpk-1(pk1393)*, and *nhr-49(nr2041);hpk-1(pk1393)* worm embryos kept in 21% O_2_ for 65 hr, and counted as ability to reach at least L4 stage (four repeats totalling >100 individual worms per strain). All comparisons not significant (ordinary one-way ANOVA corrected for multiple comparisons using the Tukey method). **(D)** The graph shows average population survival of wild-type, *hif-1(ia4)*, *hpk-1(pk1393)*, and *hif-1(ia4);hpk-1(pk1393)* worm embryos kept in 21% O_2_ for 65 hr, and counted as ability to reach at least L4 stage (four repeats totalling >60 individual worms per strain). All comparisons not significant (ordinary one-way ANOVA corrected for multiple comparisons using the Tukey method). n.s. = not significant, WT = wild-type.

**Supplementary Figure 7.**
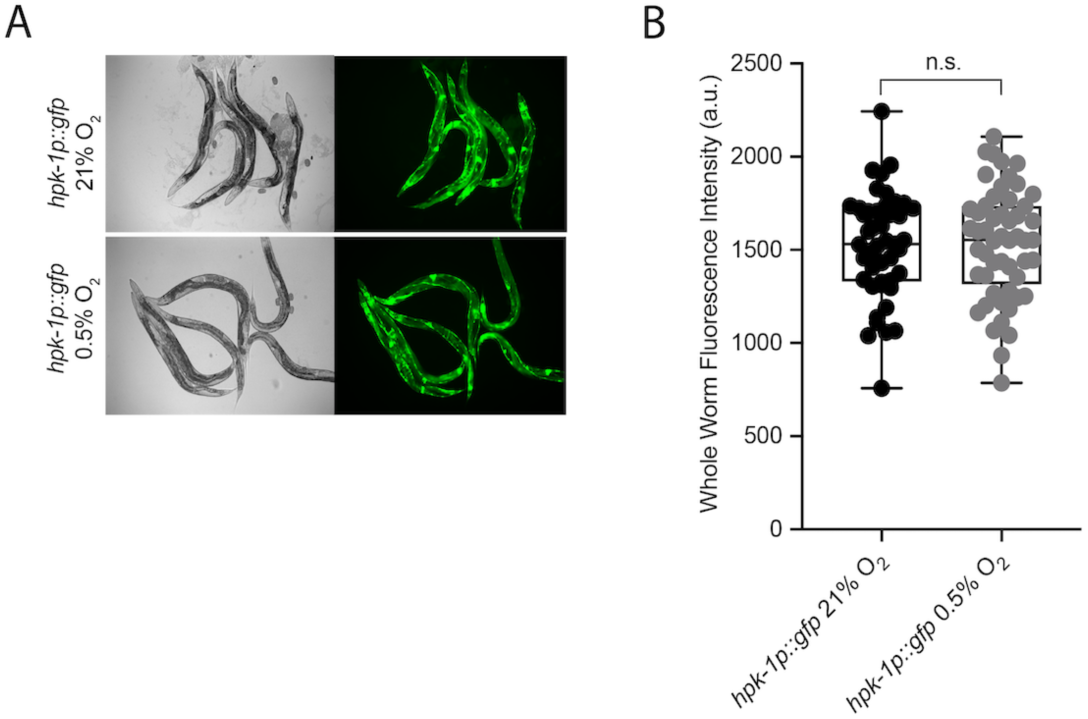
*hpk-1* is post-transcriptionally regulated in hypoxia. **(A)** Representative micrographs show *Phpk-1::gfp* adult worms in 21% O_2_ or following 4 hr exposure to 0.5% O_2_ and 1 hr recovery in 21% O_2_. **(B)** Quantification of whole worm GFP levels in *Phpk-1::gfp* worms following 4 hr exposure to 0.5% O_2_ and 1 hr recovery in 21% O_2_ or kept at 21% O_2_ (four repeats totalling >30 individual worms per strain; ordinary one-way ANOVA corrected for multiple comparisons using the Tukey method). n.s. = not significant.

## Supplementary Tables

**Supplementary Table 1.** Statistical comparison of each genotype’s ability to reach at least L4 following 24 hr exposure to 0.5% O_2_ as embryo and then allowed to recover at 21% O_2_ for 65 hr, compared to worm embryos kept in 21% O_2_ for 65 hr.

| 0.5% O<sub>2</sub> Figure | 21% O<sub>2</sub> Figure | Genotype | p-value |
| --- | --- | --- | --- |
| Figure 2A | Supplementary Figure 1A | WT | 0.4943 |
| Figure 2A | Supplementary Figure 1A | *nhr-49(nr2041)* | <0.0001**** |
| Figure 2A | Supplementary Figure 1A | *hif-1(ia4)* | <0.0001**** |
| Figure 2A | Supplementary Figure 1A | *nhr-49(nr2041);hif-1(ia4)* | <0.0001**** |
| Figure 3D | Supplementary Figure 4A | WT | 0.1403 |
| Figure 3D | Supplementary Figure 4A | *nhr-49(nr2041)* | <0.0001**** |
| Figure 3D | Supplementary Figure 4A | *fmo-2(ok2147)* | 0.0005*** |
| Figure 3D | Supplementary Figure 4A | *acs-2(ok2457)* | 0.0008*** |
| Figure 3D | Supplementary Figure 4A | *fmo-2(ok2147);acs-2(ok2157)* | <0.0001**** |
| Figure 3E | Supplementary Figure 4B | WT;*EV(RNAi)* | 0.9995 |
| Figure 3E | Supplementary Figure 4B | *nhr-49(nr2041);EV(RNAi)* | 0.0003*** |
| Figure 3E | Supplementary Figure 4B | WT;*nhr-49(RNAi)* | 0.0001*** |
| Figure 3E | Supplementary Figure 4B | *nhr-49(nr2041);nhr-49(RNAi)* | 0.0002*** |
| Figure 3E | Supplementary Figure 4B | WT;*atg-10(RNAi)* | 0.0001*** |
| Figure 3E | Supplementary Figure 4B | *nhr-49(nr2041);atg-10(RNAi)* | 0.0021** |
| Figure 3E | Supplementary Figure 4B | WT;*atg-7(RNAi)* | 0.0003*** |
| Figure 3E | Supplementary Figure 4B | *nhr-49(nr2041);atg-7(RNAi)* | 0.0534 |
| Figure 3E | Supplementary Figure 4B | WT;*bec-1(RNAi)* | 0.0015** |
| Figure 3E | Supplementary Figure 4B | *nhr-49(nr2041);bec-1(RNAi)* | 0.0033** |
| Figure 3E | Supplementary Figure 4B | WT;*epg-3(RNAi)* | 0.0006*** |
| Figure 3E | Supplementary Figure 4B | *nhr-49(nr2041);epg-3(RNAi)* | 0.0001*** |
| Supplementary Figure 5D | Supplementary Figure 5E | *EV(RNAi)* | 0.0072** |
| Supplementary Figure 5D | Supplementary Figure 5E | *nhr-49(RNAi)* | 0.0001*** |
| Supplementary Figure 5D | Supplementary Figure 5E | *nhr-67(RNAi)* | 0.0002*** |
| Figure 6F | Supplementary Figure 6C | WT | 0.9119 |
| Figure 6F | Supplementary Figure 6C | *nhr-49(nr2041)* | <0.0001**** |
| Figure 6F | Supplementary Figure 6C | *hpk-1(pk1393)* | <0.0001**** |
| Figure 6F | Supplementary Figure 6C | *nhr-49(nr2041);hpk-1(pk1393)* | <0.0001**** |
| Figure 6G | Supplementary Figure 6D | WT | 0.0223* |
| Figure 6G | Supplementary Figure 6D | *hif-1(ia4)* | <0.0001**** |
| Figure 6G | Supplementary Figure 6D | *hpk-1(pk1393)* | <0.0001**** |
| Figure 6G | Supplementary Figure 6D | *hif-1(ia4);hpk-1(pk1393)* | <0.0001**** |

All p-values are derived using ordinary one-way ANOVA corrected for multiple comparisons using the Tukey method. \*p<0.05, \*\*p<0.01, \*\*\*p<0.001, and \*\*\*\*p<0.0001. WT = wild-type.

**Supplementary Table 2.** Statistical comparison of each genotype’s ability to reach at least L4 stage from L1 stage following 48 hr exposure to 0.5% O_2_ as embryos, compared to worms kept in 21% O_2_ for 48 hr.

| 0.5% O<sub>2</sub> Figure | 21% O<sub>2</sub> Figure | Genotype | p-value |
| --- | --- | --- | --- |
| Figure 2B | Supplementary Figure 1B | WT | >0.9999 |
| Figure 2B | Supplementary Figure 1B | *nhr-49(nr2041)* | 0.0028** |
| Figure 2B | Supplementary Figure 1B | *hif-1(ia4)* | 0.0021** |
| Figure 2B | Supplementary Figure 1B | *nhr-49(nr2041);hif-1(ia4)* | <0.0001**** |

All p-values are derived using ordinary one-way ANOVA corrected for multiple comparisons using the Tukey method. \*\*p<0.01 and \*\*\*\*p<0.0001. WT = wild-type.

**Supplementary Table 3.** List of the 83 genes upregulated more than two-fold in 21% O_2_ vs. 0.5% O_2_ in wild-type and *hif-1(ia4)* worms, but not in *nhr-49(nr2041)* worms, i.e. *nhr-49*-dependent, *hif-1*-independent genes.

|  |  |  |  |  |  |  |
| --- | --- | --- | --- | --- | --- | --- |
| ABHD-5.1 | C42D4.1 | CYP-37B1* | FAAH-2 | MNK-1 | R10D12.6 | TBC-14 |
| ACS-2 | C49G7.12 | CYP-43A1* | FBXA-98 | NHL-3 | SIAH-1 | UGT-2* |
| AKT-2 | C49G7.7 | EEED8.2 | FBXA-99 | NHR-131 | SRH-2 | UGT-20* |
| ATG-2 | C50F7.5 | EPG-9 | FMO-2 | NHR-238 | SRR-6 | UGT-51* |
| B0228.6 | CBP-3 | F13E9.15 | GBA-2 | NHR-65 | SRT-39 | VEM-1 |
| B0403.3 | CUP-16 | F16B12.4 | ICL-1 | NHR-88 | SRW-86 | W09G12.7 |
| C01B4.7 | CYP-13A11* | F16C3.2 | K09D9.1 | NUMR-2 | SYX-2 | Y19D10A.4 |
| C06E4.6 | CYP-13A5* | F20B6.7 | K09E9.1 | OAC-14 | T04H1.2 | Y38C1AA.6 |
| C18B12.4 | CYP-13A6* | F22F7.4 | LGC-1 | OAC-6 | T16G1.4 | Y43F8C.3 |
| C25F9.11 | CYP-25A3* | F35E8.2 | LGG-2 | PALS-14 | T21B4.21 | Y77E11A.2 |
| C33A11.2 | CYP-34A9* | F43G6.8 | M01A8.1 | R03H10.6 | T22C8.6 | ZIP-5 |
| C33A12.3 | CYP-35A1* | F59C6.16 | MFB-1 | R09D1.12 | T24E12.5 |  |

\* genes involved in detoxification response

**Supplementary Table 4.** List of 139 genes upregulated more than two-fold in 21% O_2_ vs. 0.5% O_2_ in wild-type and *nhr-49(nr2041)* worms, but not in *hif-1(ia4)* worms, i.e. *hif-1*-dependent, *nhr-49*-independent genes.

|  |  |  |  |  |  |  |
| --- | --- | --- | --- | --- | --- | --- |
| ACS-12 | C52E2.5 | F07C3.9 | FBXA-50 | MMAA-1 | SQRD-1 | UGT-5* |
| B0507.6 | C56E6.2 | F13H8.11 | FMO-4 | NAS-28 | SRD-35 | UGT-50* |
| BATH-36 | CBP-2 | F16H6.10 | FRM-10 | NHR-161 | SRH-283 | W07A12.4 |
| C06G3.6 | CEEH-1 | F17C11.11 | GBH-2 | NHR-173 | SRX-12 | Y102A11A.9 |
| C08B6.2 | CHIL-13 | F22B3.7 | GCL-1 | NHR-195 | SRX-125 | Y105C5B.25 |
| C10C5.5 | CLEC-144 | F22B5.4 | GLB-1 | NHR-210 | SRX-21 | Y116A8C.25 |
| C14B9.3 | CLEC-222 | F25E5.4 | GLB-15 | NHR-42 | T04A11.1 | Y17G7B.8 |
| C15B12.8 | CLEC-223 | F29C6.1 | HGO-1 | NHR-59 | T07G12.5 | Y32B12C.1 |
| C18H9.5 | COMT-4 | F35E12.9 | K04C1.3 | NLG-1 | T20D4.3 | Y40H7A.11 |
| C25F9.5 | CYP-13A3* | F37H8.2 | K05C4.9 | OAC-31 | T24A6.7 | Y43F8B.13 |
| C31H5.5 | CYP-14A2* | F42C5.4 | K08B4.7 | OAC-54 | T28F3.5 | Y43F8B.15 |
| C32D5.12 | D1054.18 | F42G2.2 | K11D12.13 | OAC-7 | T28H10.1 | Y43F8B.23 |
| C32E8.9 | DDO-1 | F45D11.14 | K11G9.1 | PCK-1 | T28H10.3 | Y47H10A.5 |
| C33D9.6 | DEL-5 | F47H4.2 | KMO-1 | PCP-2 | TBC-6 | Y4C6B.4 |
| C34D1.4 | DH11.2 | F53C3.4 | LINC-72 | PGP-7 | TPRA-1 | Y53G8B.2 |
| C34D10.2 | E02C12.10 | F56D2.5 | M01H9.2 | PHY-2 | TPS-2 | Y57A10A.14 |
| C37C3.10 | ECH-9 | F57B9.1 | M03A1.3 | R05G6.10 | TWK-31 | Y5H2B.1 |
| C44C1.6 | EFK-1 | FBXA-188 | MADF-10 | R07E4.1 | UGT-17* | ZK228.4 |
| C44E12.1 | EGL-9 | FBXA-25 | MATH-27 | R08D7.7 | UGT-24* | ZK550.6 |
| C49C3.15 | ETHE-1 | FBXA-26 | MCE-1 | SKR-5 | UGT-4* |  |

\* genes involved in detoxification response

**Supplementary Table 5.** List of 264 genes upregulated more than two-fold in 21% O_2_ vs. 0.5% O_2_ via RNA-seq in wild-type, *nhr-49(nr2041)*, and *hif-1(ia4)*.

|  |  |  |  |  |  |  |
| --- | --- | --- | --- | --- | --- | --- |
| AAKG-4 | CATP-3 | F21D12.3 | FBXA-82 | M01H9.3 | SLC-17.4 | UGT-19* |
| ARRD-11 | CDR-2 | F22H10.2 | FBXA-91 | M163.1 | SLC-36.5 | UGT-33* |
| ARRD-24 | CHIL-18 | F25B3.5 | FBXA-92 | M60.7 | SODH-1 | UGT-54* |
| ARRD-8 | CKR-2 | F26F12.3 | FBXL-1 | MAI-1 | SQST-1 | W04C9.8 |
| B0205.13 | CNC-2 | F27D9.2 | FIPR-22 | MTL-1 | SRD-27 | W05H9.1 |
| B0205.14 | CNC-4 | F28H1.1 | FIPR-24 | NEP-26 | SRH-48 | Y15E3A.5 |
| B0310.3 | CNG-1 | F28H6.8 | FIPR-26 | NHR-103 | SRI-36 | Y34F4.4 |
| B0462.5 | COEL-1 | F33H12.7 | FKH-7 | NHR-107 | SRI-39 | Y34F4.7 |
| BEST-5 | COMT-5 | F34H10.3 | FTN-1 | NHR-115 | SRM-3 | Y37A1B.5 |
| BIGR-1 | CYP-13A8* | F37A8.5 | GEM-4 | NHR-126 | SRP-8 | Y38H6C.9 |
| C02F5.12 | CYP-14A1* | F40F12.9 | GLO-3 | NHR-132 | STO-1 | Y39A3A.4 |
| C04A11.5 | CYP-14A4* | F41E6.5 | GPA-1 | NHR-18 | STR-31 | Y42G9A.1 |
| C04C11.25 | CYP-14A5 | F43C1.7 | H28G03.1 | NHR-211 | SWT-1 | Y43F8B.14 |
| C06B3.6 | CYP-32B1* | F43C11.7 | HAF-7 | NHR-212 | T05H4.15 | Y43F8B.9 |
| C06B3.7 | CYP-33C7 | F43H9.4 | HIL-1 | NHR-226 | T08A9.13 | Y44A6C.1 |
| C06E1.11 | CYP-33C8* | F45D3.4 | HPD-1 | NHR-228 | T09F5.12 | Y45F10D.6 |
| C08E8.4 | D1086.5 | F45E1.5 | HRG-1 | NHR-230 | T10C6.15 | Y46G5A.36 |
| C08F11.13 | DAAO-1 | F46A8.13 | HRG-2 | NHR-57 | T10G3.1 | Y47G6A.5 |
| C10C5.2 | DC2.5 | F47B10.9 | HSP-12.3 | NHR-79 | T10H9.8 | Y47H10A.3 |
| C11G10.1 | DCT-1 | F47B8.3 | HSP-70 | NHR-90 | T12A7.6 | Y54G11A.7 |
| C18A11.1 | DCT-7 | F47B8.4 | IRG-2 | NHR-99 | T12D8.5 | Y54G2A.11 |
| C23H4.2 | DCT-8 | F53A9.7 | IST-1 | NIPI-3 | T16G1.5 | Y54G2A.36 |
| C23H4.6 | DOD-3 | F53B2.8 | K01C8.1 | NNT-1 | T19C4.5 | Y54G2A.52 |
| C24B5.4 | E03H4.8 | F53C3.6 | K01F9.2 | PALS-34 | T20D4.7 | Y56A3A.33 |
| C25F9.12 | E04F6.6 | F54B8.4 | K02D7.1 | PALS-6 | T24C4.4 | Y58A7A.3 |
| C28G1.6 | EGAP1.1 | F55C12.19 | K05B2.4 | PARG-2 | T27F6.8 | Y58A7A.4 |
| C29F7.2 | F08G12.5 | F56C4.4 | K06A9.2 | PCS-1 | T28B8.1 | Y58A7A.5 |
| C31H2.4 | F09F7.6 | F56D6.8 | K06G5.3 | PEK-1 | T28F4.4 | Y6E2A.4 |
| C33D9.13 | F10E9.12 | F57B9.3 | K08D8.12 | PGP-3 | T28F4.5 | Y6G8.2 |
| C35A5.6 | F13C5.1 | F58G6.9 | K08D9.4 | PGP-9 | TLI-1 | Y71G12B.2 |
| C36B1.6 | F13D11.3 | F59B10.4 | K09C8.7 | PITR-5 | TOS-1 | ZC239.14 |
| C37A5.3 | F14F9.2 | F59E11.7 | K10G4.3 | PTR-22 | TSP-1 | ZC395.5 |
| C44H9.5 | F14F9.3 | FBXA-105 | K10G6.9 | R06B9.5 | TTR-23 | ZC443.3 |
| C46C2.2 | F14F9.4 | FBXA-163 | K12H6.6 | R06C1.6 | TTR-37 | ZC443.4 |
| C49G7.10 | F15A8.6 | FBXA-189 | KGB-2 | R08F11.4 | TTS-1 | ZIG-7 |
| C54F6.18 | F15E6.3 | FBXA-24 | KREG-1 | R102.1 | UBC-8 | ZK836.3 |
| C54G10.1 | F17C11.13 | FBXA-66 | LINC-37 | R186.1 | UGT-13* |  |
| CAH-4 | F18G5.6 | FBXA-80 | M01B2.13 | SCL-2 | UGT-18* |  |

\* genes involved in detoxification response

**Supplementary Table 6.** Worm strains used in this study.

| Strain | Genotype | Reference |
| --- | --- | --- |
| N2 | Wild-type | (Brenner, 1974) |
| STE68 | *nhr-49(nr2041) I* | (Marc R. Van Gilst et al., 2005) |
| VE40 | *eavEx20[Pfmo-2::*gfp* + *rol-6(su1006)*]* | (Goh et al., 2018) |
| STE129 | *nhr-49(nr2041) I; eavEx20[Pfmo-2::*gfp* + *rol-6(su1006)*]* | This study |
| ZG31 | *hif-1(ia4) V* | (Jiang et al., 2001) |
| STE130 | *nhr-49(nr2041) I; hif-1(ia4) V* | This study |
| VC1668 | *fmo-2(ok2147) IV* | (Leiser et al., 2015) |
| RB1899 | *acs-2(ok2457) V* | (J. Zhang et al., 2011) |
| STE131 | *fmo-2(ok2147) IV; acs-2(ok2457) V* | This study |
| STE110 | *nhr-49let13) I* | (K. Lee et al., 2016) |
| AGP33a | *nhr-49(nr2041)I; glmEx5 [Pnhr-49::nhr-49::gfp + Pmyo-2::mCherry]* | (Naim et al., 2020) |
| AGP65 | *nhr-49(nr2041)I; glmEx9 [Pgly-19::nhr-49::gfp + Pmyo-2::mCherry]* | (Naim et al., 2020) |
| AGP53 | *nhr-49(nr2041)I; glmEx11 [Pcol-12::nhr-49::gfp + Pmyo-2::mCherry]* | (Naim et al., 2020) |
| AGP51 | *nhr-49(nr2041)I; glmEx13 [Prgef-1::nhr-49::gfp + Pmyo-2::mCherry]* | (Naim et al., 2020) |
| WBM170 | *wbmEx57 [Pacs-2::gfp + rol-6(su1006)]* | (Burkewitz et al., 2015) |
| WBM169 | *nhr-49(nr2041) I; wbmEx57 [Pacs-2::gfp + rol-6(su1006)]* | (Burkewitz et al., 2015) |
| AGP25f | *glmEx5 (Pnhr-49::nhr-49::gfp + Pmyo-2::mCherry)* | (Ratnappan et al., 2014) |
| EK273 | *hpk-1(pk1393) X* | (Raich et al., 2003) |
| STE132 | *nhr-49(nr2041) I; hpk-1(pk1393) X* | This study |
| STE133 | *hif-1(ia4) V; hpk-1(pk1393) X* | This study |
| STE117 | *nhr-49(et13) I; eavEx20[Pfmo-2::gfp + rol-6(su1006)]* | (Goh et al., 2018) |
| AVS394 | *artEx12 [Phpk-1::gfp + rol-6(su1006)]* | (Das et al., 2017) |

**Supplementary Table 7.**
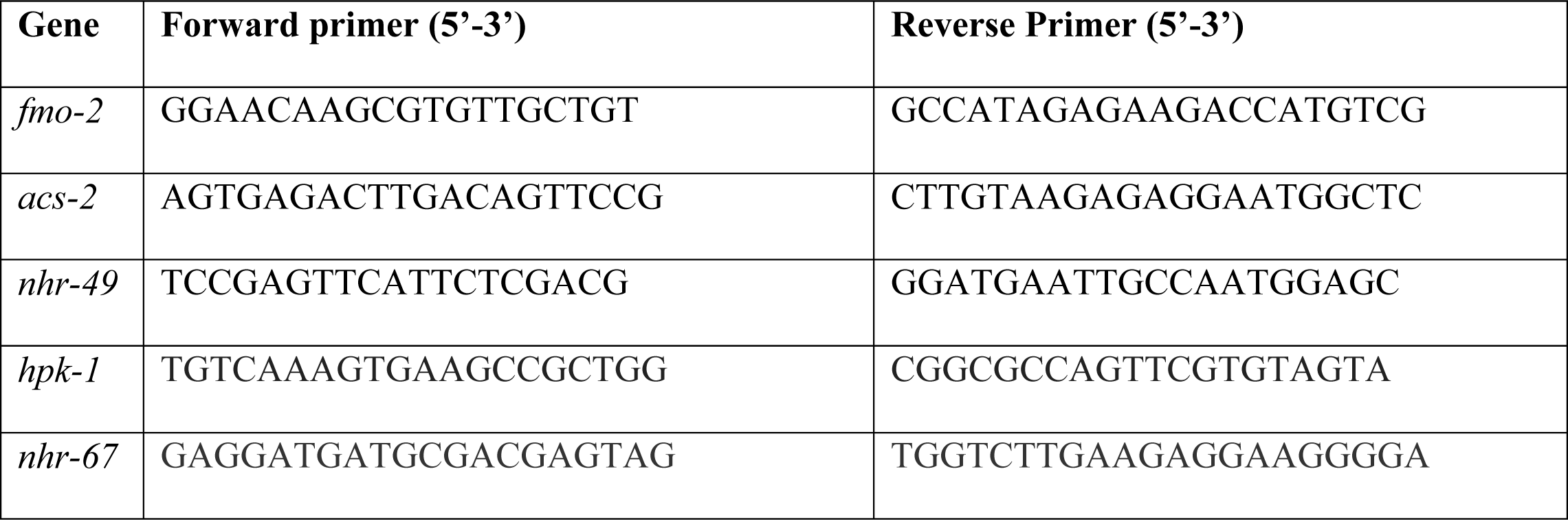
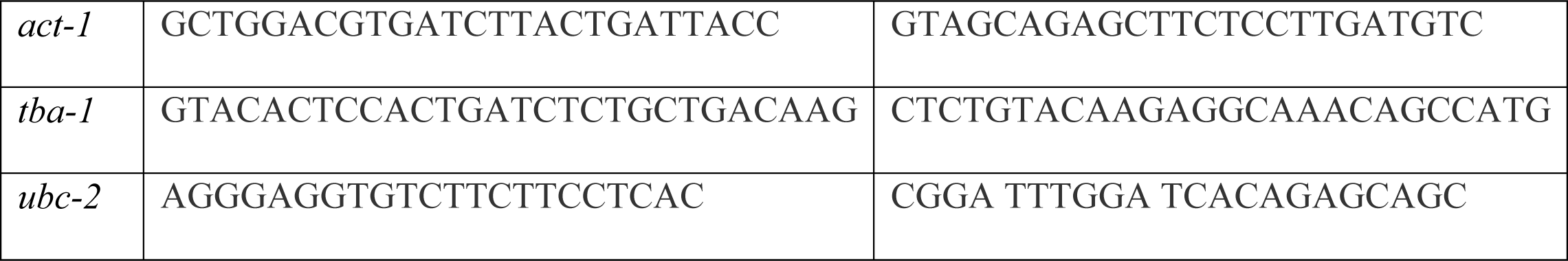
List of qRT-PCR primer sequences used in this study.

